# Protective effect of pre-existing natural immunity in a nonhuman primate reinfection model of congenital cytomegalovirus infection

**DOI:** 10.1101/2023.04.10.536057

**Authors:** Matilda Moström, Shan Yu, Dollnovan Tran, Frances Saccoccio, Cyril J. Versoza, Daniel Malouli, Anne Mirza, Sarah Valencia, Margaret Gilbert, Robert Blair, Scott Hansen, Peter Barry, Klaus Früh, Jeffrey D. Jensen, Susanne P. Pfeifer, Timothy F. Kowalik, Sallie R. Permar, Amitinder Kaur

**Affiliations:** Tulane National Primate Research Center, Tulane University, Covington LA; Duke Human Vaccine Institute, Duke University, Durham, NC; Center for Evolution & Medicine, School of Life Sciences, Arizona State University, Tempe, AZ; Oregon Health and Sciences University, Beaverton, OR; University of Massachusetts Chan Medical School, Worcester, MA; University of California, Davis, CA; Weill Cornell Medicine, New York, NY

## Abstract

Congenital cytomegalovirus (cCMV) is the leading infectious cause of neurologic defects in newborns with particularly severe sequelae in the setting of primary CMV infection in the first trimester of pregnancy. The majority of cCMV cases worldwide occur after non-primary infection in CMV-seropositive women; yet the extent to which pre-existing natural CMV-specific immunity protects against CMV reinfection or reactivation during pregnancy remains ill-defined. We previously reported on a novel nonhuman primate model of cCMV in rhesus macaques where 100% placental transmission and 83% fetal loss were seen in CD4^+^ T lymphocyte-depleted rhesus CMV (RhCMV)-seronegative dams after primary RhCMV infection. To investigate the protective effect of preconception maternal immunity, we performed reinfection studies in CD4+ T lymphocyte-depleted RhCMV-seropositive dams inoculated in late first / early second trimester gestation with RhCMV strains 180.92 (*n*=2), or RhCMV UCD52 and FL-RhCMVΔRh13.1/SIV*gag*, a wild-type-like RhCMV clone with SIV*gag* inserted as an immunological marker (*n*=3). An early transient increase in circulating monocytes followed by boosting of the pre-existing RhCMV-specific CD8+ T lymphocyte and antibody response was observed in the reinfected dams but not in control CD4+ T lymphocyte-depleted dams. Emergence of SIV Gag-specific CD8+ T lymphocyte responses in macaques inoculated with the FL-RhCMVΔRh13.1/SIV*gag* virus confirmed reinfection. Placental transmission was detected in only one of five reinfected dams and there were no adverse fetal sequelae. Viral whole genome, short-read, deep sequencing analysis confirmed transmission of both reinfection RhCMV strains across the placenta with ∼30% corresponding to FL-RhCMVΔRh13.1/SIV*gag* and ∼70% to RhCMV UCD52, consistent with the mixed human CMV infections reported in infants with cCMV. Our data showing reduced placental transmission and absence of fetal loss after non-primary as opposed to primary infection in CD4+ T lymphocyte-depleted dams indicates that preconception maternal CMV-specific CD8+ T lymphocyte and/or humoral immunity can protect against cCMV infection.

**Author Summary:** Globally, pregnancies in CMV-seropositive women account for the majority of cases of congenital CMV infection but the immune responses needed for protection against placental transmission in mothers with non-primary infection remains unknown. Recently, we developed a nonhuman primate model of primary rhesus CMV (RhCMV) infection in which placental transmission and fetal loss occurred in RhCMV-seronegative CD4+ T lymphocyte-depleted macaques. By conducting similar studies in RhCMV-seropositive dams, we demonstrated the protective effect of pre-existing natural CMV-specific CD8+ T lymphocytes and humoral immunity against congenital CMV after reinfection. A 5-fold reduction in congenital transmission and complete protection against fetal loss was observed in dams with pre-existing immunity compared to primary CMV in this model. Our study is the first formal demonstration in a relevant model of human congenital CMV that natural pre-existing CMV-specific maternal immunity can limit congenital CMV transmission and its sequelae. The nonhuman primate model of non-primary congenital CMV will be especially relevant to studying immune requirements of a maternal vaccine for women in high CMV seroprevalence areas at risk of repeated CMV reinfections during pregnancy.

## Introduction

Human cytomegalovirus (CMV) is a betaherpesvirus that results in lifelong persistent infection. While infection in immunocompetent hosts is typically asymptomatic, CMV causes life-threatening illness in immunosuppressed hosts such as transplant recipients and individuals with untreated HIV infection. CMV is also the most common cause of congenital infection in newborns with severe neurological sequelae resulting from primary infection in CMV-seronegative women in the first trimester of pregnancy (1). Congenital CMV (cCMV) can also follow non-primary infection in pregnant CMV-seropositive women. Non-primary infection due to either reactivation of endogenous latent CMV or reinfection with diverse CMV strains is the most common cause of human cCMV cases worldwide occurring in regions with high CMV seroprevalence rates among women of reproductive age (2, 3, 4, 5, 6). Even in the United States, three-quarters of cCMV infections reported between 1988-1994 were attributed to non-primary infection (7). Consequently, the development of a maternal vaccine to prevent cCMV has been a tier 1 priority of the National Institute of Medicine for two decades but has faced several challenges (8). A major barrier to vaccine development is that the immune determinants of protection against vertical transmission, particularly after non-primary infection, remain elusive. Even though natural CMV infection induces a robust and durable humoral and cellular immune response, pre-existing immunity is not sufficient to prevent reinfections or cCMV. High maternal seroprevalence regions in the world can account for >500,000 cCMV infant births per year despite “low” transmission rates of 1-2% To have worldwide relevance, an effective CMV vaccine will need to prevent both primary and non-primary cCMV. Insight in to the role of pre-existing natural immunity in limiting congenital CMV transmission and its protective components are thus of paramount importance for the rational design of an effective maternal vaccine against cCMV.

Several studies have reported a reduced fetal infection rate after non-primary compared to primary CMV infection. A meta-analysis of epidemiologic studies conducted between 1966 and 2006 revealed that the rate of intra-uterine CMV transmission after primary infection during pregnancy was 32%, as opposed to 1.4% after non-primary infection In a prospective study of 2,378 woman/newborn pairs with maternal CMV serology at first visit and CMV screening of newborns at birth, the risk of maternal-fetal transmission after primary infection was 4-fold higher compared to non-primary infection (11). A similar reduction in transmission after non-primary infection has been reported from cCMV cohorts in Italy and Brazil where maternal primary and non-primary infections were monitored by serial serology (12, 13). It is worth noting that in contrast to transmission, the symptomatology and severity of cCMV with respect to sensorineural hearing loss and neurodevelopmental anomalies can be comparable in non-primary and primary infection (14, 15, 16). These data suggest that the mechanisms protecting against vertical transmission may be different from those determining the outcome of fetal infection once transmission has occurred. Elucidating the determinants of protection against acquisition and disease severity, particularly the role of pre-conceptional immunity in CMV-seropositive pregnant women, is of paramount importance for the development of a maternal vaccine effective against both primary and non-primary cCMV infection (13, 17). In this regard, the rhesus macaque CMV reinfection pregnancy model has the potential to address some of the unanswered questions pertaining to non-primary cCMV, including the rate of placental transmission, protective effects of pre-existent natural immunity, and the host response to reinfection.

The rhesus macaque animal model offers several translational benefits for studying the pathogenesis and immunology of human CMV (HCMV) infection. The rhesus CMV (RhCMV) genome is closely related to HCMV and the natural history and biology of RhCMV infection in rhesus macaques bears important similarities to HCMV infection (18, 19, 20). Natural RhCMV infection is widespread in colony-bred rhesus macaques; near 100% seroprevalence rates are reached within a year of birth with natural acquisition occurring as a result of horizontal transmission from close contact with infectious body fluids containing shed virus (21, 22, 23). Thus, breeding rhesus macaques have been CMV-seropositive for at least 2-3 years before reaching the age of sexual maturity. This mimics the natural history and infection profile of HCMV in regions of the world where CMV-seroprevalence in women of child-bearing age exceeds 95% as a result of primary infections occurring in childhood (17). The rhesus macaque model of RhCMV infection is thus well suited for investigating immune determinants of protection against non-primary cCMV infection. We previously reported on a placental transmission model of primary cCMV infection in rhesus macaques (24). In this model, we observed 100% vertical transmission and 83% fetal loss in CD4+ T lymphocyte-depleted CMV-seronegative macaques infected with RhCMV in late first / early second trimester gestation, with protection conferred by passive infusion of high potency anti-CMV antibodies prior to infection (24, 25). Here we report the first instance of cCMV in CMV-seropositive rhesus macaques, representing a nonhuman primate (NHP) non-primary cCMV model.

Similar to our primary infection studies, pregnant macaques at late first / early second trimester gestation were subjected to CD4+ T lymphocyte depletion and then inoculated with RhCMV virus strains. The difference here was that the experiments were performed in CMV-seropositive macaques with pre-existing naturally acquired RhCMV-specific immunity that were then experimentally inoculated with RhCMV to simulate reinfection during pregnancy. In a subset of animals we used FL-RhCMVΔRh13.1/SIV*gag*, a wild-type-like RhCMV clone with *SIVgag* inserted as an immunological marker (26), for experimental reinfection to enable *in vivo* tracking of the reinfection virus and distinguish it from reactivation of endogenous RhCMV. In contrast to primary RhCMV infection, reinfection of CD4+ T lymphocyte-depleted RhCMV-seropositive dams resulted in reduced placental transmission and complete protection against fetal loss, accompanied by a robust innate immune response and boosting of pre-existent CMV-specific immunity. Our study demonstrates the feasibility of using a NHP animal model to study non-primary cCMV infection and demonstrates the protective effect of pre-existing natural CMV-specific CD8+ T lymphocytes and humoral immunity in a biologically relevant model of human cCMV infection.

## Results

### Reinfection model of congenital CMV in rhesus macaques

To investigate the protective effect of natural pre-existing CMV-specific immunity against placental transmission, we applied the study design of our previously published CD4+ T lymphocyte depletion primary cCMV infection model (24). We repeated the study design but substituted primary infection studies in RhCMV-seronegative macaques with non-primary reinfection studies in CD4+ T lymphocyte-depleted RhCMV-seropositive macaques with naturally acquired RhCMV infection. Colony-bred rhesus macaques have high RhCMV seroprevalence rates with natural acquisition of CMV by one year of age and thus have established pre-existing RhCMV-specific immunity for at least two years before reaching sexual maturity at 3-4 years of age (21, 23). The reinfection experiments were performed after CD4+ T lymphocyte depletion using the same dose and timeframe of infection as in the primary infection model which had yielded 100% placental transmission. Hence, comparison of placental transmission in CD4+ T lymphocyte-depleted macaques with primary RhCMV infection and RhCMV reinfection enabled an evaluation of the protective effect of pre-existing natural RhCMV-specific CD8+ T lymphocyte and humoral immunity in RhCMV-seropositive dams.

Five RhCMV-seropositive dams were enrolled in the “RhCMV-seropositive Reinfection” group and three RhCMV-seropositive dams were enrolled in the “RhCMV-seropositive Control” group to control for endogenous RhCMV reactivation without reinfection (**Fig 1A**, **Table 1**). All dams were enrolled in the first trimester of pregnancy and subjected to *in vivo* CD4+ T lymphocyte depletion at late first trimester / early second trimester between 49 to 59 gestation days, similar to the historical control group of CD4+ T lymphocyte depletion and primary RhCMV infection (**Table 1**). One week after administration of the CD4+ T lymphocyte depleting antibody, the reinfection group dams were inoculated with different RhCMV strains. Two RhCMV-seropositive dams received an intravenous inoculation of the fibroblast-passaged UL128-complex intact RhCMV strain 180.92 (27, 28) at 2×10^6^ TCID_50_ and were followed until natural birth at 152 and 164 gestational days, respectively (**Table 1**, **Fig 1A**). Three dams were intravenously inoculated with 1×10^6^ pfu each of RhCMV strains UCD52 and full-length recombinant RhCMV expressing SIV*gag* (FL-RhCMVΔRh13.1/SIV*gag*), and followed up until an elective Cesarian section (C-section) at mean 144 (range 142-147) days gestation (**Fig 1A**, **Table 1**). The choice of RhCMV 180.92 and RhCMV UCD52 virus strains were based on previously used isolates shown to cross the placenta in primary infection studies (24, 25, 29). FL-RhCMVΔRh13.1/SIV*gag*, a full-length RhCMV generated by repairing strain 68-1 with wild-type sequence that includes a heterologous SIV*gag* sequence as a foreign gene in the place of Rh13.1 (26), was used for reinfection experiments in three macaques. The full length clone enabled distinction of endogenous virus from the exogenous challenge inoculum in RhCMV-seropositive animals and allowed confirmation of reinfection as previously demonstrated in RhCMV/SIV vaccine studies (30, 31, 32). Three RhCMV-seropositive dams that underwent CD4+ T lymphocyte depletion without reinfection were included as a control group to monitor placental transmission due to reactivation of endogenous RhCMV virus strains (**Table 1**, **Fig 1A**).

**Fig 1.**
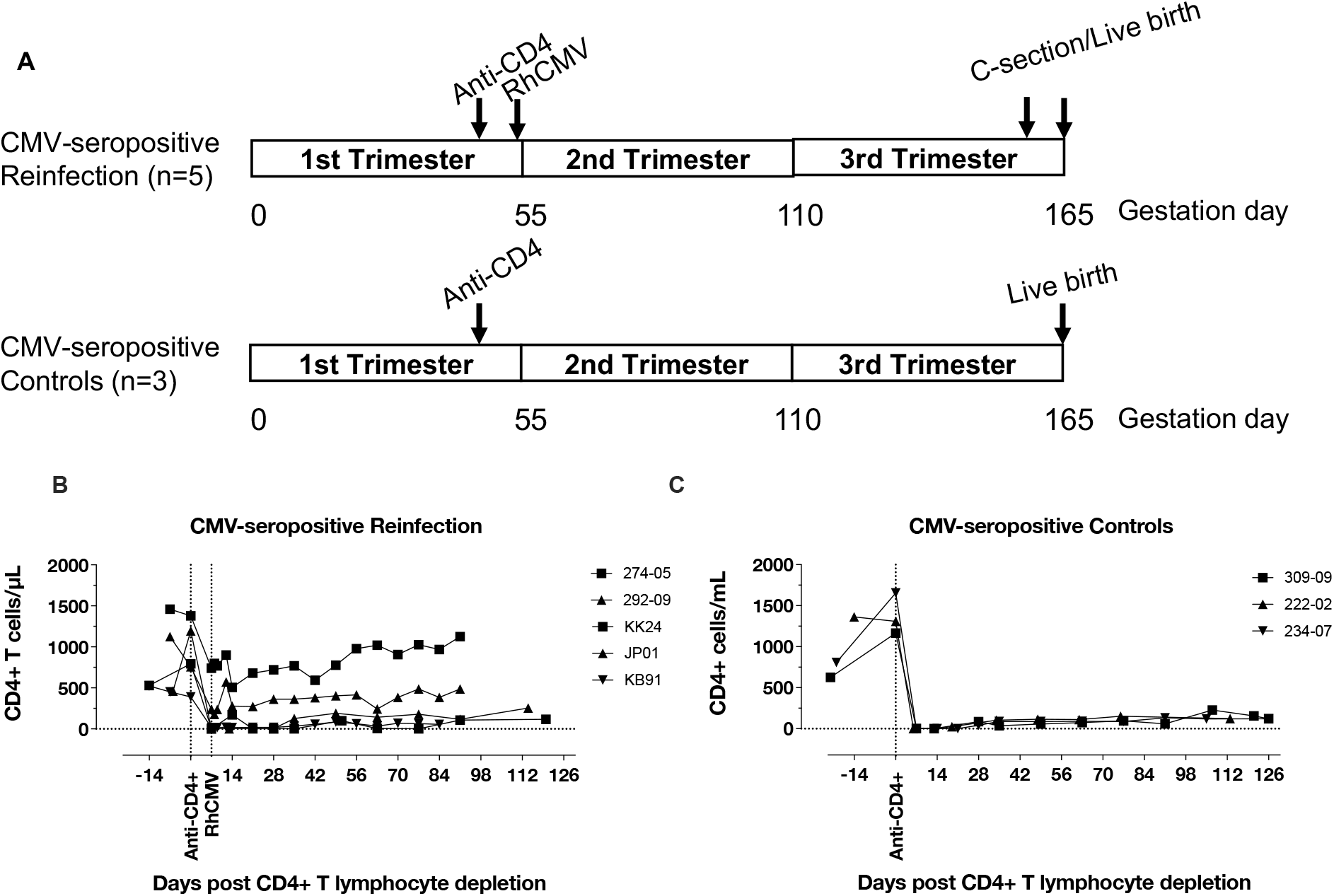
Study design and kinetics of CD4+ T lymphocyte depletion in experimental groups. (A) Schematic of study design of cCMV transmission in pregnant CMV-seropositive rhesus macaques. (B-C) Peripheral blood CD4+ T lymphocyte counts following anti-CD4 antibody administration in (B) CMV-seropositive Reinfection group (*n*=5); and (C) CMV-seropositive Control group (*n*=3).

**Table 1.** Study outline of animal groups.

|  | Number of animals |  | RhCMV Inoculum | Age in years<br>Mean (Range) | Gestational day<br>Mean (Range) |  |  |
| --- | --- | --- | --- | --- | --- | --- | --- |
|  |  |  |  |  | CD4+ T lymphocyte depletion | RhCMV inoculation | Study termination |
| CMV-seropositive Reinfection | n=5 | n=2 | • RhCMV 180.92 | 5.5 (4-9) | 53.2 (49-59) | 60.2 (56-66) | 144 (142-147) |
|  |  | n=3 | • RhCMV UCD52<br>• FL-RhCMVΔRh13.1/SIVgag |  |  |  |  |
| CMV-seropositive Controls | n=3 | n=3 | Not inoculated | 7.0 (4-11) | 52.0 (50-55) | Not inoculated | 164 (155-171) |
| CMV-seronegative Primary Infection* | n=6 | n=1 | • RhCMV 180.92 | 11.6 (4-16) | 50.5 (47-56) | 57.5 (54-63) | 94 (74-165) |
|  |  | n=5 | • UCD52<br>• UCD59<br>• RhCMV 180.92 |  |  |  |  |

\*Bialas, et al. PNAS (2015). Nelson, et al. JCI Insight (2017)

#### CD4+ T lymphocyte and RhCMV viral dynamics

Following administration of a single dose of 50mg/kg rhesus recombinant anti-CD4 monoclonal antibody (clone CD4R1), profound CD4 depletion, resulting in loss of >95% of circulating CD4+ T lymphocytes, was observed at day 7 in three out of five RhCMV-seropositive reinfected animals as well as in all three RhCMV-seropositive controls (**Fig 1B-C**). The peripheral blood CD4+ T lymphocyte count in these animals declined from a mean baseline value of 950 cells/μL to less than 65 cells/μL throughout the study period. Two RhCMV-seropositive reinfected macaques (JP01 and KK24) showed relatively suboptimal depletion with persistent circulating CD4+ T lymphocytes at 46-68% of baseline values (**Fig 1B**).

RhCMV DNA was monitored in the plasma, saliva, urine and amniotic fluid of reinfection and control dams (**Fig 2A-B**). Immunocompetent RhCMV-seropositive macaques are typically aviremic but shed RhCMV in body fluids such as urine and saliva (21). Following CD4+ T lymphocyte depletion, low levels of plasma RhCMV DNA were detected intermittently in two out of five RhCMV-seropositive reinfection dams (KK24 and KB91) but in none of the three CMV-seropositive control dams (**Fig 2A-B**). Placental RhCMV transmission as evidenced by detection of RhCMV DNA in the amniotic fluid was detected in one of five RhCMV-seropositive reinfected dams (KK24) at day 7 post RhCMV inoculation (**Fig 2A**) but not in any of the CD4+ T lymphocyte depletion control animals (**Fig 2B**). Consistent with previous reports, virus shedding as evidenced by detection of RhCMV DNA in the saliva and urine was present in all eight RhCMV-seropositive dams at baseline (21). A one-log or greater increase in virus shedding in the saliva and / or urine was detected in all five reinfected dams at 1-9 weeks post reinfection (**Fig 2A**). It was also seen in all three control dams post CD4+ T lymphocyte depletion (**Fig 2B**) suggesting that increased viral shedding could result from both reactivation and reinfection. Taken together, these data demonstrate placental transmission after RhCMV reinfection but not after reactivation. The transmitter dam KK24 had low-level transient viremia post reinfection, however this did not coincide temporally with the time point of detection of RhCMV DNA in the amniotic fluid (**Fig 2B**).

**Fig 2.**
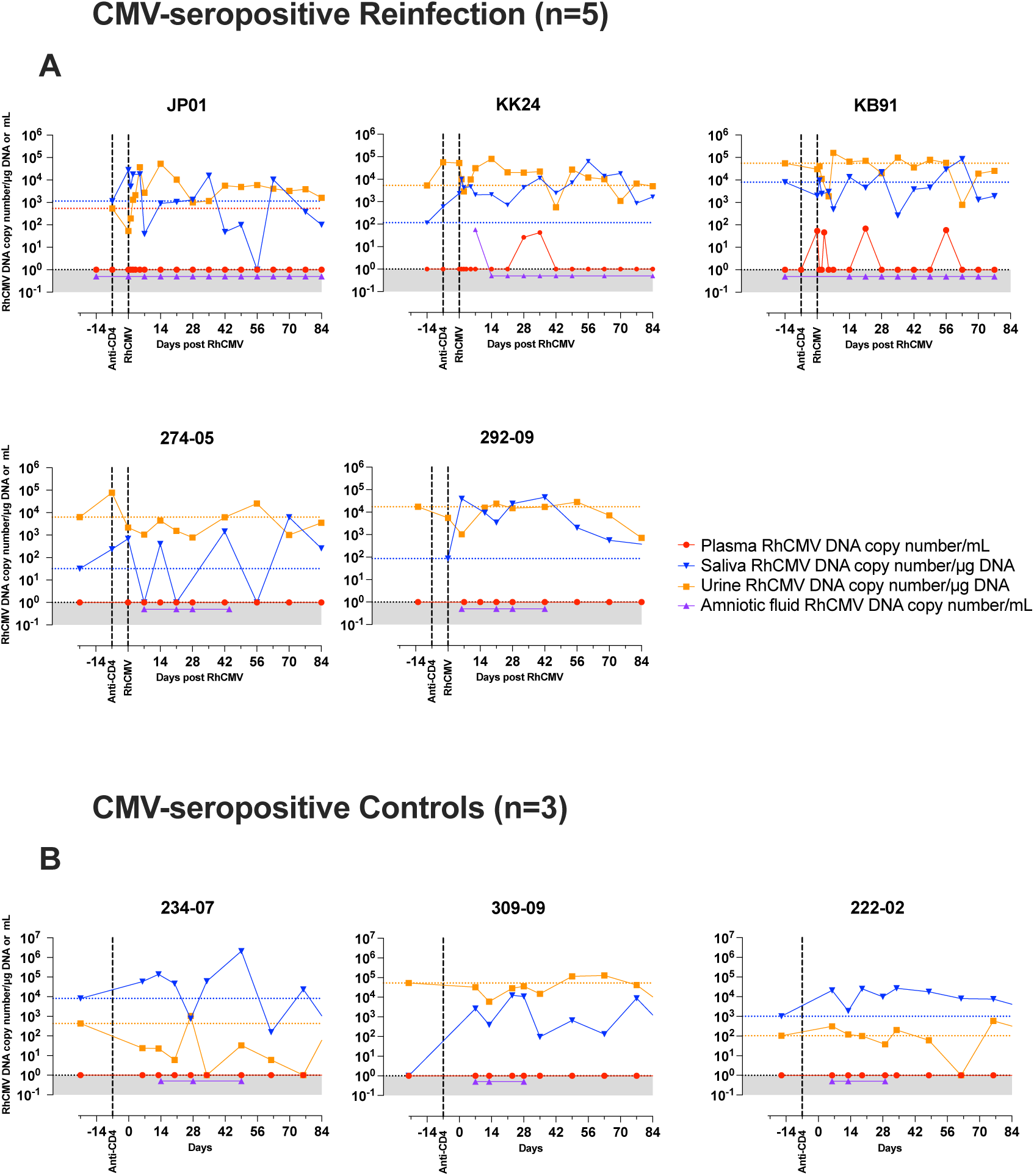
RhCMV viral kinetics in blood and body fluids of individual CD4+ T lymphocyte depeleted CMV-seropositive macaques. RhCMV in plasma (indicated in red), saliva (blue), urine (orange), and amniotic fluid (purple) in (A) five CMV-seropositive reinfected animals and (B) three CMV-seropositive control animals. Plasma and amniotic fluid are reported in RhCMV DNA copy number/mL of sample fluid; saliva and urine are reported as RhCMV DNA copy number/μg of input DNA. In CMV-seropositive controls, the equivalent post-infection time-points on the x-axis are aligned concurrently with the CMV-seropositive Reinfection group. The black vertical lines indicate time of anti-CD4 antibody (CD4R1) and RhCMV inoculation. Animals JP01, KK24, and KB91 were inoculated with RhCMV UCD52 and FL-RhCMVΔRh13.1/SIV*gag*; animals 274-05 and 292-09 were inoculated with RhCMV 180.92; animals 234-07, 309-09, and 222-02 remained without a reinfection. The horizontal stippled line indicates the baseline RhCMV DNA copy number/μg of input DNA in either saliva (blue) or urine (orange).

### Innate and adaptive host response to RhCMV reinfection

Several features of innate immune activation were evident within one week of RhCMV reinfection. A 3-to 100-fold non-significant (p-value=0.25; non-parametric Wilcoxon Signed Rank test) increase in frequency of circulating HLA-DR^hi^CD14^+^ monocytes from 3.1 ± 3.7% (mean ± SD) to 14.3 ± 10.9% was observed at day 1-5 post reinfection in RhCMV-seropositive pregnant macaques (**Fig 3**, **Fig S1A**). Concurrently, there was also an expansion of lineage (Lin)-HLA-DR^hi^ dendritic cells, Lin-HLA-DR-SSC^hi^CD14^+^ putative myeloid-derived suppressor cells (MDSC) and a transient decline in circulating B lymphocytes which did not reach statistical significance (p-value=0.25; non-parametric Wilcoxon Signed Rank test) (**Fig 3**, **Fig S1A-C)**. There was also no significant change in the frequency of circulating natural killer (NK) cells and CD8+ T lymphocytes (**Fig 3**). Evidence of innate immune activation was corroborated by an increase in plasma analytes, notably IL-8 and MIP-1b, as measured by Luminex in the first week post reinfection (**Fig S2A**). Of note, an increase in eotaxin was only observed in KK24, the transmitter dam (**Fig S2A**). Overall, these data indicate a rapid activation of the innate immune system predominantly involving myeloid cells in response to RhCMV reinfection.

**Fig 3.**
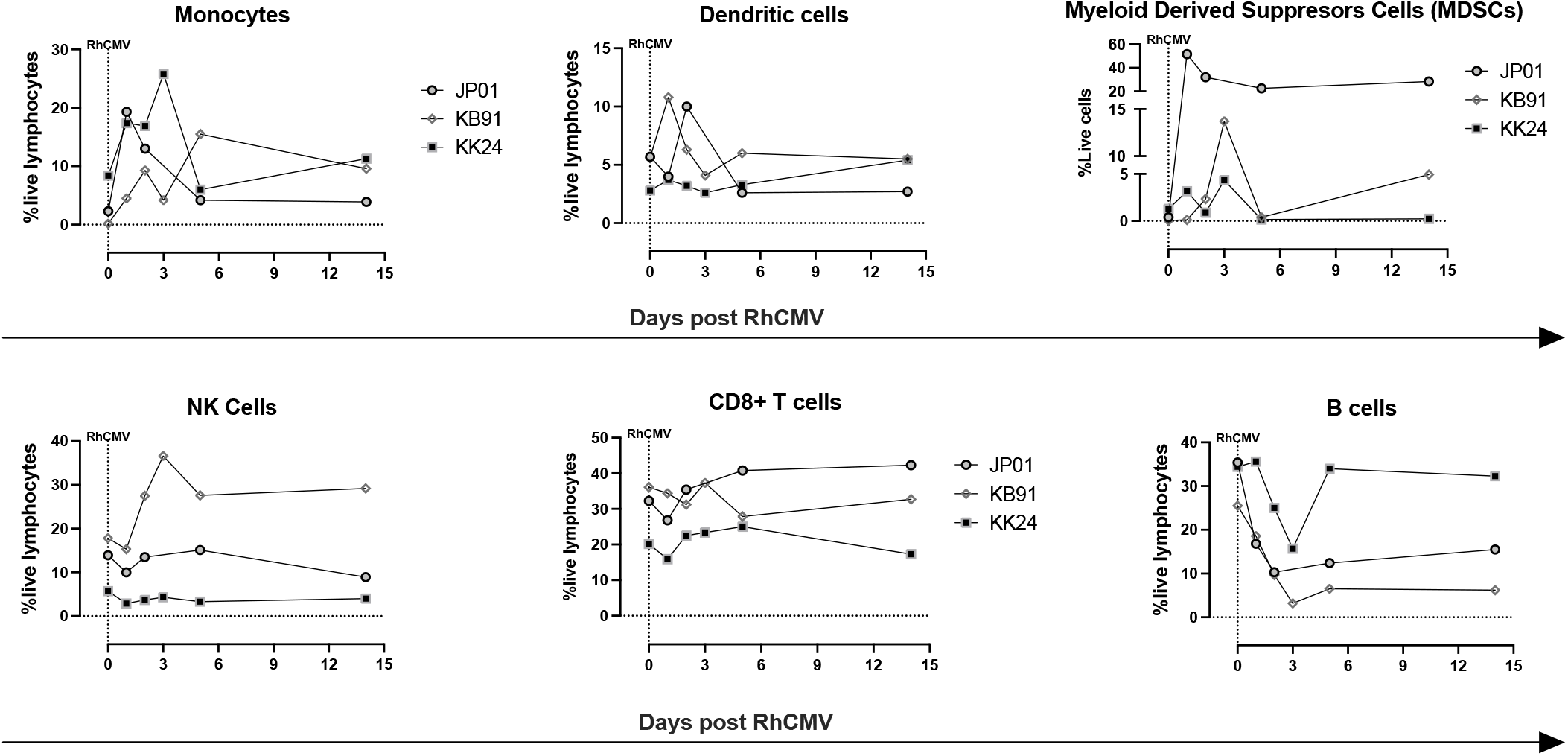
Early immunophenotypic changes following RhCMV reinfection in CMV-seropositive macaques. Immunophenotyping of circulating peripheral blood mononuclear cells in acute RhCMV reinfection. Plots show the kinetics of different lymphocyte subsets in three CMV-seropositive reinfected macaques (JP01, KB91, and KK24). Paired t-test comparing baseline prereinfection values with values at time-point of maximal change in the first 7 days post reinfection was performed.

To evaluate the effect of RhCMV reinfection on pre-existing adaptive immunity, we longitudinally monitored RhCMV-specific CD8+ T lymphocyte responses and anti-RhCMV antibodies in the CD4+ T lymphocyte-depleted reinfection and control macaques. Memory RhCMV-specific CD8+ T lymphocyte responses against the RhCMV immediate early 1 (IE1), IE2 or pp65 peptide pools measured by intracellular cytokine staining assay were detected at baseline in all the CMV-seropositive macaques. RhCMV-specific CD8+ T lymphocyte responses were also measured at 8-10 weeks post reinfection and compared with baseline responses. Four dams in the RhCMV-seropositive reinfection group had baseline immunodominant CD8+ T lymphocyte responses targeting the IE1 or IE2 peptide pools which increased 2-to 10-fold following reinfection indicating a booster effect (**Fig 4**). An increase in CD107a expression, and IFN-γ, IL-2, and TNF-α cytokine secretion was observed post reinfection with a significant increase in the frequency of TNF-α cytokine secreting RhCMV-specific CD8+ memory T lymphocytes (**Fig 4A**). Boolean analysis of all four effector functions showed a trend for decreased proportion of monofunctional and increased proportion of polyfunctional RhCMV-specific CD8+ T cell responses post reinfection (*P-*value 0.11; **Fig 4B**). This was associated with a significant increase in the frequency of RhCMV-specific 4-functional (*P-*value 0.03) and dual CD107a+IFN-γ+ positive (*P-*value 0.03) CD8+ T lymphocytes post reinfection compared to pre-reinfection values (**Fig 4C**). Analysis of pre-reinfection RhCMV-specific CD8+ T lymphocyte responses in individual animals revealed that the transmitter dam KK24 had the the lowest frequency (0.91%) as well as the highest proportion of monofunctional responses at baseline compared to the other reinfection dams (**Fig S2B**). However, the post reinfection boost in this frequency was comparable in the transmitter and non-transmitter dams (**Fig S2B**). It is noteworthy that the increase in magnitude and polyfunctionality of memory RhCMV-specific CD8+ T lymphocytes post reinfection occurred despite the presence of CD4+ T lymphocyte depletion.

**Fig 4.**
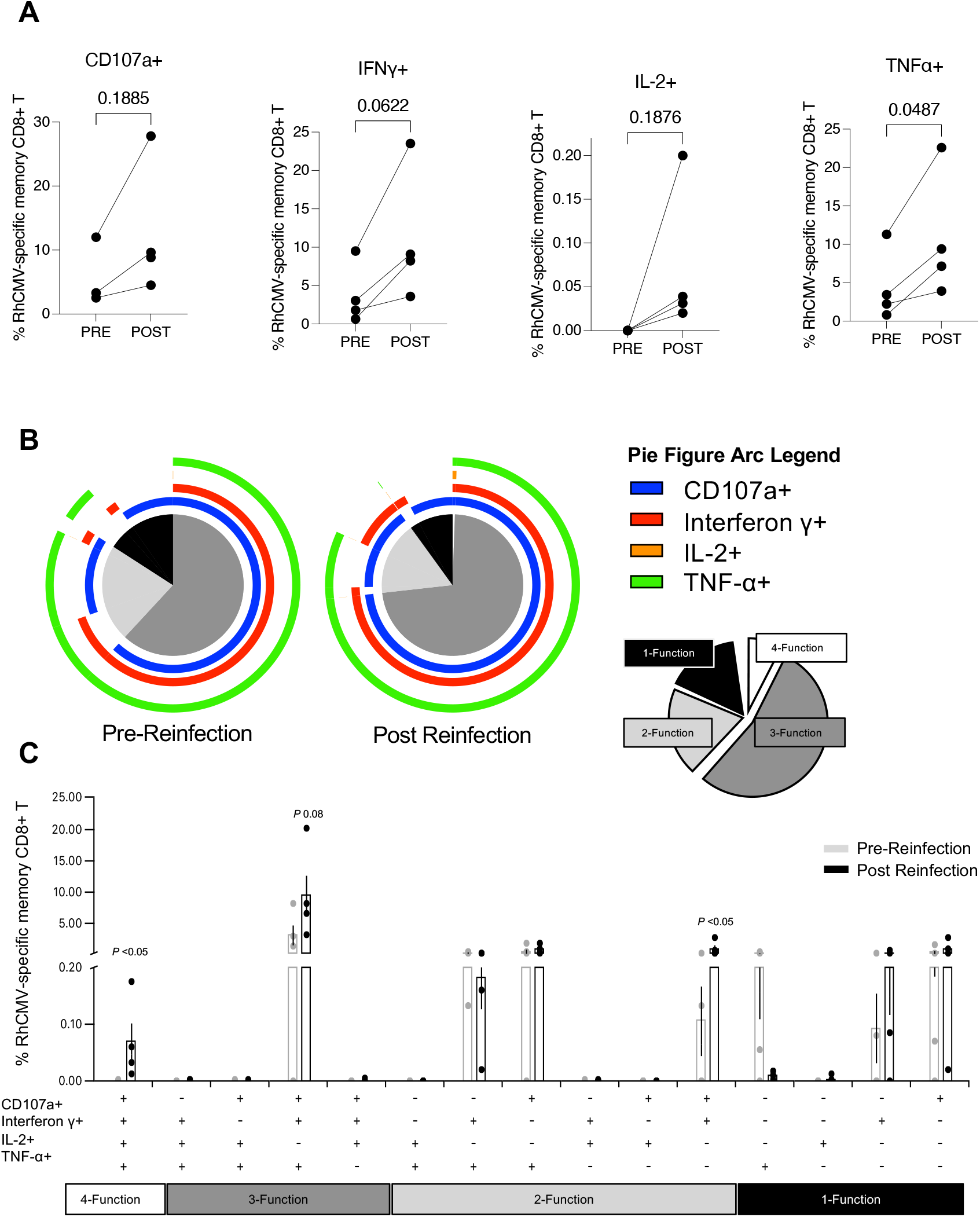
CMV-specific CD8+ T lymphocyte memory responses to RhCMV immediate early (IE) proteins and exogenous SIV Gag protein in CD4+ T lymphocyte depleted RhCMV reinfected macaques. (A) Paired IE-specific responses by CD107a expression and secretion of IFN-γ, IL-2, and TNF-α in CMV-seropositive macaques (n=4) reinfected with RhCMV UCD52 and FL-RhCMVΔRh13.1/SIV*gag*. Pre-reinfection responses were compared with responses at week 8-10 post RhCMV reinfection using paired t-test. (B) Polyfunctional SPICE analysis of IE-specific responses pre vs post RhCMV reinfection. CD107a (blue arc), IFN-γ (red arc), IL-2 (orange arc), and TNF-α (green arc). Four-functional responses are displayed in white, three-functional responses in dark grey, two-functional response in light grey, and mono-functional responses in black. (C) Bar graph of the polyfunctional responses pre (grey) and post (black) RhCMV reinfection (*n*=4). The RhCMV IE-specific response was measured by intracellular cytokine staining after stimulation with RhCMV IE1 and / or IE2 peptide pools depending on the baseline immunodominant response in individual animals. Comparison of pre-and post reinfection Boolean responses were compared with the Wilcoxon rank sum test using SPICE v6 software.

We also monitored end-point dilution titers of anti-RhCMV gB-binding IgG responses and fibroblast neutralization activity against RhCMV 180.92 (**Fig 5**). Following reinfection, a transient 0.8-to >1.0-log increase in anti-gB binding titers was observed in three RhCMV-reinfected macaques (**Fig. 5A**). Analysis of individual animals showed the two macaques (KK24 and JP01) that experienced a suboptimal CD4+ lymphocyte depletion (46-68% loss) responded with a 1.0-log or greater increase in magnitude of anti-gB IgG responses (**Fig 5A**), suggesting that CD4+ T lymphocyte help may have aided a boost of anti-gB antibodies against pre-existing and new specifities post RhCMV reinfection. In a group analysis, baseline values for gB-binding and neutralizing antibody levels at pre-CD4+ T lymphocyte depletion time-points were comparable between the RhCMV-seropositive reinfected and RhCMV-seropositive control dams (**Fig 5B**). Post CD4+ T lymphocyte depletion, a change in antibody titers was only observed in the reinfection dams with significant increases in the first 8 weeks post reinfection compared to the RhCMV-seropositive control dams (**Fig. 5C-D**).

**Fig 5.**
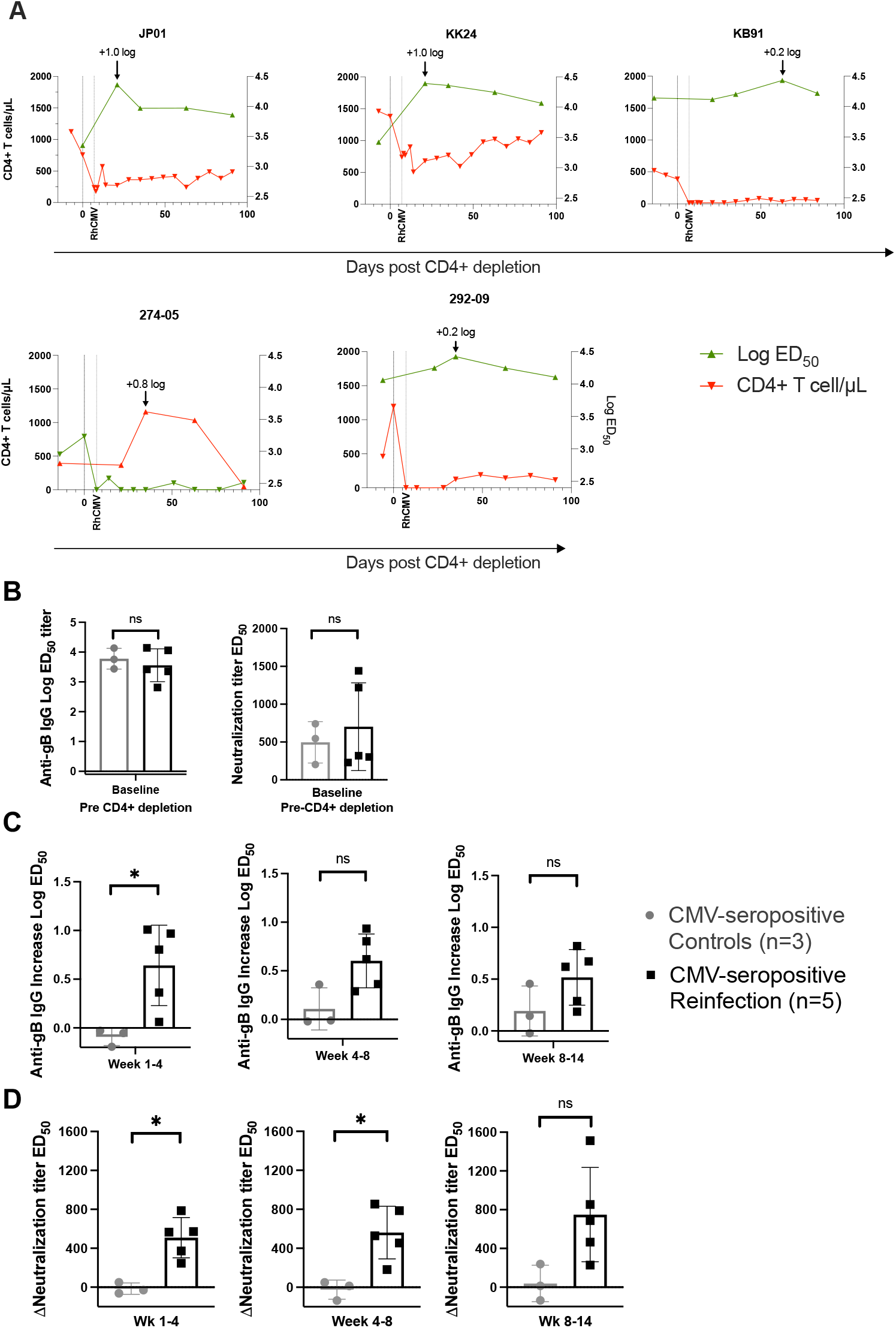
RhCMV-specific antibody responses in CD4+ T lymphocyte depleted CMV-seropositive reinfected macaques. (A) Kinetic data of anti-gB binding antibodies in graphs on individual animals showing log ED_50_ of anti-gB binding titer (green line) superimposed on CD4+ lymphocytes/μL (red line). (B) Comparison of CMV-seropositive reinfected dams (*n*=5) and CMV-seropositive controls (*n*=3) by their anti-gB binding titer and fibroblast neutralization against RhCMV 180.92 at baseline preceding CD4+ lymphocyte depletion. (C) Maximal increase in baseline anti-gB ED_50_ compared between the two groups. (D) Comparison of maximal neutralization titer change from baseline over time in both CMV-seropositive controls and CMV-seropositive reinfected dams. ED_50_ = Effective Dose 50

In contrast to the reinfected animals, the CD4+ T lymphocyte depleted CMV-seropositive control macaques showed no change in the RhCMV-specific CD8+ memory response post CD4+ T lymphocyte depletion (**Fig S3A**). Neither did they display an increase in anti-gB IgG levels post CD4+ T lymphpcyte depletion (**Fig S3B**). In all, our results demonstrate elevation of RhCMV-specific CD8+ T lymphocyte and antibody responses that was evident only after RhCMV reinfection, not reactivation following CD4+ T lymphocyte depletion.

### Placental transmission of reinfection virus

Although the boosting of pre-existing RhCMV-specific immune responses and the increase in viral shedding were suggestive of reinfection, natural fluctuations of endogenous virus replication could also lead to changes in shed virus. To determine if reinfection virus was being shed, we evaluated the three macaques (KB91, KK24, and JP01) that were infected with the clone FL-RhCMVΔRh13.1/SIV*gag* virus containing the exogenous transgene *SIVgag*. Screening the saliva and urine at every time-point in each of the three macaques revealed a low positive signal by SIV*gag*-specific real time PCR in the saliva of one animal, KB91, at a single time-point (**Fig 6A**). However, SIV Gag-specific IFN-γ-secreting memory CD8+ T lymphocyte responses were detected in all three macaques at 6 weeks or later post reinfection and ranged in frequency between 0.9 to 1.4% of circulating memory CD8+ T lymphocytes (**Fig 6B**). An antibody assay was also developed to determine if these animals generated antibody responses to the SIV Gag protein. No measureable responses were found in the dams which may be due to a near absence of viremia in the reinfected animals (**Fig S4**). Cumulatively, these data provide evidence for successful RhCMV reinfection of the CD4+ T lymphocyte-depleted CMV-seropositive dams.

**Fig. 6.**
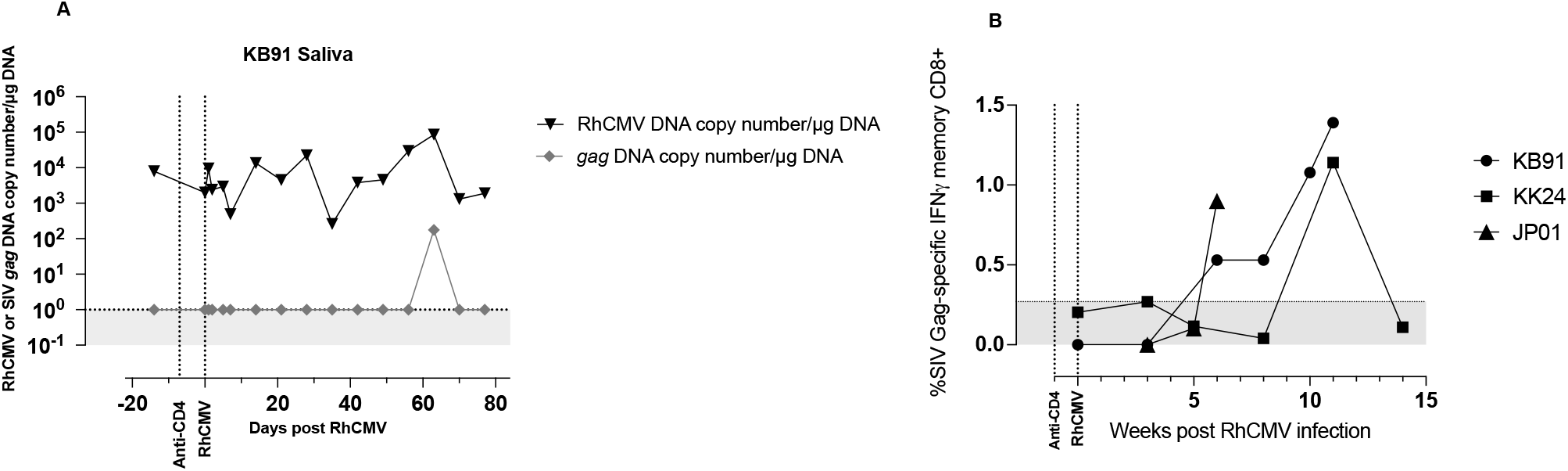
Evidence of reinfection in CD4+ T lymphocyte depleted FL RhCMVΔRh13.1/SIV*gag* inoculated dams. (A) RhCMV-specific (black line) and SIV*gag*-specific (grey line) real time PCR results in the saliva of one CMV-seropositive reinfected animal (KB91). All other animals were found negative for SIV*gag* DNA in saliva and urine. (B) Detection of SIV Gag-specific T lymphocyte responses measured longitudinally against a SIVmac239 Gag peptide pool in CMV-seropositive reinfected macaques (KB91, KK24, and JP01) inoculated with RhCMV UCD52 and FL-RhCMVΔRh13.1/SIV*gag*. Horizontal stippled line shows negative cut-off based on pre-reinfection values.

To confirm the passage of reinfection RhCMV virus across the placenta, amniotic fluid DNA from the transmitter dam KK24 was amplified by multiple displacement amplification (MDA) and viral DNA enriched by PCR-amplification using RhCMV-specific primer pairs. Amplicons were then sequenced to >10,000X coverage on a Ion Torrent Sequencer. The resulting whole genome sequencing data (mean read length: ∼200 bp) was mapped against the RhCMV UCD52 and FL-RhCMVΔRh13.1/SIV*gag* reference assemblies. Investigations of uniquely mapping regions demonstrated placental transmission of both reinfection RhCMV strains with ∼70% corresponding to RhCMV UCD52 and ∼30% to FL-RhCMVΔRh13.1/SIV*gag* consistent with the mixed human CMV infections reported in infants with cCMV (33).

### Protective effect of natural pre-existing immunity against congenital CMV transmission

To evaluate protection conferred by pre-conception immunity, we compared viral and pregnancy outcome parameters in the CD4+ T lymphocyte-depleted RhCMV-seropositive reinfection macaques with the CD4+ T lymphocyte-depleted primary RhCMV infection historical control animals (**Fig 7 and Table 2**). The CD4+ T lymphocyte-depleted RhCMV-seropositive reinfected dams showed intermittent, detectable RhCMV DNAemia which was significantly lower compared to RhCMV-seronegative animals with primary infection (**Fig 7A**). The CD4+ T lymphocyte-depleted RhCMV-seropositive controls showed no detectable RhCMV DNAemia (**Fig 7A**). Amniotic fluid sampled at weekly intervals showed variable detection of RhCMV transmission across the study groups as shown in a heatmap (**Fig 7B**). While all six dams in the primary infection group had detectable amniotic fluid RhCMV DNA (mean range 49-580 copies/ml) at one or more sampling time-points, amniotic fluid RhCMV DNA at mean±S.D 57±146 copies/mL was detected in only one animal in the reinfection group at a single time-point (**Fig 7B**). Amniotic fluid RhCMV transmission was not detected in any of the CD4+ T lymphocyte depletion RhCMV-seropositive controls (**Fig 7B**). The fetuses of the mothers with natural pre-existing immunity fared better with 100% fetal survial (**Fig 7C-D**). Overall, placental transmission was reduced from 100% in primary infection to 20% in the reinfection group (*P* <0.05; Log-rank test), whereas fetal survival was increased from 16% to 100% in the reinfection groups (*P* <0.05; Log-rank test) (**Table 2**, **Fig 7C-D**). RhCMV DNA PCR evaluation of placental tissues and fetal tissues confirmed a protective role of maternal pre-existing immunity in preventing the spread and replication of RhCMV *in vivo* during first trimester infection of rhesus macaques (**S1 Table**). The fetal growth parameters in the CD4+ T lymphocyte-depleted RhCMV reinfected dams were comparable to the RhCMV-seropositive control dams (**S5 Fig**) and within the normal range of reference values established in rhesus macaques (34).

**Fig 7.**
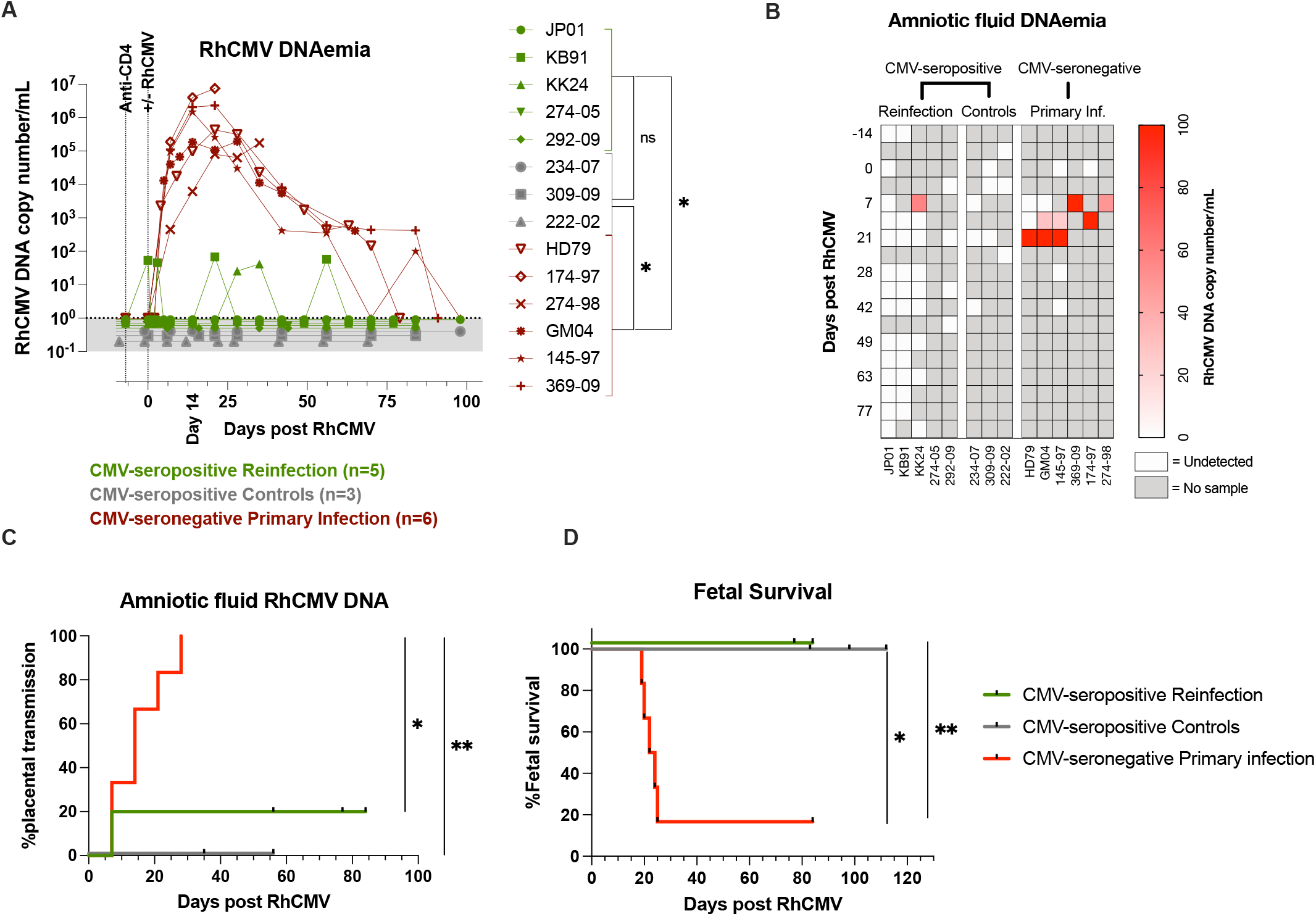
Protective effect of pre-existing immunity against congenital CMV transmission. (A) Plasma RhCMV-specific PCR in CD4+ T lymphocyte depleted CMV-seronegative primary infected macaques (red; *n*=6) compared to both CMV-seropositive reinfected (green; *n*=5) and CMV-seropositive controls (grey; *n*=3). Area Under the Curve (AUC) values of plasma RhCMV DNA between 0-99 days were compared between groups using the Man-Whitney test. P-values <0.05 denoted with a single * were considered significant. (B) Heatmap of RhCMV-specific DNA copy number in amniotic fluid in CMV-seronegative primary infected macaques (*n*=6), CMV-seropositive reinfected (*n*=5), and CMV-seropositive controls (*n*=3). (C) Kaplan-Meir curve showing cCMV frequency based on RhCMV DNA detection in the amniotic fluid in CMV-seronegative primary infected macaques (*n*=6), CMV-seropositive reinfected (*n*=5), and CMV-seropositive controls (*n*=3). (D) Kaplan-Meir curve showing fetal survival in CMV-seronegative primary infected macaques (*n*=6), CMV-seropositive reinfected (*n*=5), and CMV-seropositive controls (*n*=3). Statistical comparisons by Log-rank (Mantel-Cox) test showing significance levels: *=<0.05 and **=<0.01.

**Table 2.** Improved study outcome in dams with pre-existing immunity. The main readouts of the study are described as a frequency of total number of animals.

| Group name | Natural Immunity to RhCMV | Amniotic fluid PCR RhCMV DNA | Placental and fetal tissue PCR RhCMV DNA | Number of animals |
| --- | --- | --- | --- | --- |
| CMV-seropositive Reinfection | Yes | 20% | 0% | 5 |
| CMV-seropositive Controls | Yes | 0% | 0% | 3 |
| CMV-seronegative Primary Infection | No | 100% | 100% | 6 |

## Discussion

In this study we provide the first evidence of placental transmission in a nonhuman primate model of non-primary cCMV and demonstrate that pre-conception maternal CMV-specific immunity from previous natural CMV infection has a protective effect against congenital transmission and disease. By conducting a longitudinal experimental superinfection study with an inoculum of defined RhCMV virus stocks administered at a known gestation time-point in pregnant RhCMV-seropositive rhesus macaque dams, we could determine the risk of vertical transmission after non-primary CMV infection and compare it with historical controls from our previous studies of primary cCMV in this animal model. Moreover, by including a RhCMV virus with a foreign transgene (SIV*gag*), we could track one of the inoculated RhCMV viruses *in vivo* and distinguish it from endogenous RhCMV in the reinfected animals. Our data show that even in the setting of CD4+ T lymphocyte depletion, RhCMV-reinfected RhCMV-seropositive dams showed only 20% placental transmission and did not suffer any adverse pregnancy outcome or fetal infection. These findings are in sharp contrast to primary RhCMV infection in CD4+ T lymphocyte-depleted CMV-seronegative rhesus macaque dams, which resulted in 100% placental transmission and 83% fetal loss (24, 25). In the absence of CD4+ T lymphocytes, the protective effect of prior natural RhCMV infection was accompanied by an initial activation of the innate immune system in the first week after reinfection followed by boosting of pre-existing memory RhCMV-specific CD8+ T lymphocyte and antibody responses. These data point to redundant pathways of immune-mediated protection and suggest that vaccine approaches harnessing different arms of the immune system will be needed to prevent cCMV infection.

RhCMV-seropositive pregnant macaques model the immunity of reproductive age women in a high CMV seroprevalence settings. The RhCMV-seropositive macaques used in this study acquired naturally circulating RhCMV strains prevalent in the primate center colony and were never experimentally infected prior to this study. Thus, their pre-existing immune status was a result of natural RhCMV infection akin to human populations susceptible to non-primary infections. As demonstrated previously, reinfection of RhCMV-seropositive animals is enabled by viral T cell evasion mechanisms (32). However, although T cells cannot prevent reinfection due to viral immune evasion there was a clear impact of pre-existing immunity on congenital infection. In the current study, we saw a 5-fold reduction in transmission rate after non-primary infection as compared to primary infection despite the presence of CD4+ T lymphocyte depletion – akin to a recent study in humans (11). However, unlike in humans, we did not find any evidence of fetal infection or pathology. Because the diagnosis of non-primary cCMV in humans is based on the screening of newborns, and because there are no data on amniotic fluid CMV load monitoring in pregnant CMV-seropositive women, we cannot exclude the possibility of placental transmission without fetal infection also occurring in human subjects. The reduction in transmission and absence of fetal infection is indicative of a protective role of natural pre-conceptional immunity. However, there are caveats to extrapolating the findings in the NHP model to non-primary cCMV in humans. The small number of animals in this study precludes definitive conclusion of the transmission rate following non-primary infection. Furthermore, the risk of placental transmission following a single reinfection event, as was the design of the current study, does not capture the potential of multiple reinfections occurring in CMV-seropositive pregnant women in high CMV seroprevalence regions when exposed to frequent high levels of CMV shedding from different sources. Annualized CMV seroconversion rates in CMV-seronegative women are directly related to the extent of exposure to CMV shedding with 20% or higher rates in high CMV seroprevalence areas (13, 35). The same pattern may hold true for reinfection rates.

The extent to which non-primary CMV infection is a result of reactivation or reinfection, or a combination of both factors, is not known. In one of the first studies to document reinfection as an important cause of non-primary cCMV, 62% of mothers with cCMV births compared to 13% with normal births developed new antibody specificities against glycoprotein H during pregnancy suggestive of reinfection (4). More recently, the contribution of reinfection to non-primary cCMV has been deduced from detection of new antibody specificities to strain-specific polymorphic regions in the glycoproteins gB and gH (5, 6). Using these criteria, the annualized rate of reinfection in CMV-seropositive women has ranged between 10-35% in different studies with reinfections being associated with an increased risk of cCMV (5, 6). In addition to reinfection, new strain-specificities could also be due to emergence of a previously subdominant or quiescent endogenous CMV due to reactivation, but this has not been established. Our model looked at both modes of non-primary CMV infection in pregnant rhesus macaques, albeit in a small number of animals. As controls for the group of CD4+ T lymphocyte-depleted reinfected macaques, we used CD4+ T lymphocyte depletion alone to model endogenous CMV reactivation without reinfection. We inferred reactivation from a one-log or greater increase in RhCMV load shed in the urine or saliva at one or more post CD4+ T lymphocyte depletion time points. Unlike adult humans where CMV shedding is detected only in a subset of individuals, group-housed RhCMV-seropositive rhesus macaques invariably have detectable RhCMV DNA shed in the urine and/or saliva (21, 36). Consistent with this observation, all the RhCMV-seropositive macaques enrolled in this study had detectable presence of RhCMV DNA shed in the urine and/or saliva at baseline prior to any intervention. A one-log or greater increase in RhCMV shedding at post CD4+ T lymphocyte depletion time-points was detected in both the reinfection and control group of animals suggestive of reactivation. However, transient viremia and amniotic fluid transmission were only detected in the RhCMV-seropositive, CD4+ T lymphocyte depleted, reinfected dams and not the RhCMV-seropositive, CD4+ T lymphocyte depleted controls. One of the viremic reinfected dams, macaque KK24, transmitted the virus across the placenta. Confirmation of reinfection was supported by the emergence of immune responses against the foreign transgene present in the inoculated RhCMV virus, and detection of this foreign gene in shed virus. These findings were confirmed by detection of the reinfection virus in the amniotic fluid of the transmitter dam by genome-wide deep sequencing. In the absence of sequencing RhCMV shed in the urine and saliva prior to reinfection we cannot ascertain if endogenous reactivated RhCMV also crossed the placenta in the transmitter dam.

Sequencing analysis confirmed that the two RhCMV strains used for reinfection had both crossed the placenta, with RhCMV UCD52 being more prevalent than FL-RhCMVΔRh13.1/SIV*gag*. Passage of the RhCMV UCD52 strain was consistent with the findings in our primary infection studies where UCD52 was the dominant virus in the circulation and amniotic fluid (24, 25). This raises the question of what might determine placental transmission of reinfection virus. Aside from protective immunity, viral diversity and inoculum size are likely to be important determinants of cCMV transmission after reinfection. Recent studies have shown the surprising diversity of naturally circulating HCMV, which can be in the range of that observed in RNA viruses such as dengue virus and HIV (37). Moreover, mixed strain infections, such as the one observed here, are common in CMV-seroimmune individuals and reinfection cCMV (33, 38, 39). The presence of significant diversity between RhCMV strains may be an important factor determining which virus strains are transmitted. For example, viral diversity of the reinfection RhCMV from the endogenous circulating strains could render it less susceptible to immune control and hence more transmissible. Based on the sequencing data, the bottleneck of transmission appears to be wide as both virus inoculums were detected in the amniotic fluid. This phenomenon is similar to our findings in primary infection in CD4+ T lymphocyte-depleted dams where the genetic composition of RhCMV in the blood and amniotic fluid after a mixed infection with three RhCMV viruses was similar and did not reveal a readily detectable transmission bottleneck even in the presence of immune pressure exerted by pre-existing passively infused antibodies (29). However, with the current data we are unable to exclude the endogenous virus from being present in the transmitting sequences. Future sequencing analysis of the endogenous vs challenge sequences are required to answer this and to determine the relative divergence between endogenous and inoculum viruses between the animals.

Despite the absence of CD4+ T lymphocytes, RhCMV-seropositive macaques showed evidence of protection against cCMV transmission and infection. The protective effect was likely mediated by both innate immunity and CMV-specific adaptive immunity. Protection in the absence of CD4+ T lymphocytes points to protection mediated by the CD8+ T lymphocyte and humoral components of adaptive immunity. It is noteworthy that despite CD4+ T lymphocyte depletion and potential lack of CD4 help, there was still a robust memory CMV-specific CD8+ T lymphocyte or the antibody response post reinfection. Boosting of CD8+ T lymphocyte and antibody responses were only seen in reinfection animals and not in the CD4+ T lymphocyte depletion control animals. The endogenous anti-gB IgG antibodies and the pre-reinfection CMV-specific CD8+ T lymphocyte responses were significantly increased following reinfection when compared to controls. The increase in gB-specific IgG and RhCMV-specific CD8+ T lymphocyte responses is unlikely to have been a result of RhCMV reactivation as it was not observed in the CD4+ T lymphocyte depleted controls without reinfection. Of note, rhesus macaques that were both CD4-and CD8 T lymphocyte-depleted preceding kidney transplant did not see a rise in anti-gB IgG titer, suggesting that it is unlikely that our observation of boosted immune responses are a consequence of reactivation (40). It is interesting that the magnitude of the increase in antibody responses was greatest in the two macaques with partial CD4+ T lymphocyte depletion suggesting a facilitatory role for CD4 help in potentiating humoral immunity. The magnitude of the boosted memory CMV-specific CD8+ T lymphocyte response however did not appear to be affected by the extent of CD4+ T lymphocyte depletion. In the absence of epitope mapping, we cannot determine whether the amplified responses were anamnestic in nature with expansion of pre-existing or cross-reactive specificities against reinfection virus or included new responses directed towards new epitopes in the reinfecting strain.

The small number of animals in this study with only one placental-transmitter dam prevents analysis of protective correlates based on differences in immunity between transmitters and non-transmitters. Observationally, the placental-transmitter dam (KK24) displayed certain features that were different from the non-transmitter dams. For example, an increase in plasma eotaxin following reinfection was only detected in the transmitter dam. KK24 also had the lowest frequency of RhCMV IE-specific CD8+ T lymphocyte responses at baseline prior to reinfection. In addition, KK24 was one out of two dams that showed transient viremia post reinfection and that had a partial CD4+ T lymphocyte depletion as well as a 1.0 log or greater increase in anti-gB IgG response following reinfection. Whether this could have resulted in an increase in low affinity binding antibodies and transmission facilitated by transcytosis of immune complexes is an interesting possibility. Studies with a larger group of animals are warranted.

In conclusion, establishment of the NHP reinfection cCMV model has provided definitive evidence for a role of pre-conceptional natural immunity in CMV-seropositive individuals to partially protect against cCMV. Strikingly, this protection is evident even in the absence of CD4+ T lymphocytes and likely involves multiple arms of the immune system including CD8+ T lymphocyte-mediated and humoral immunity. Importantly, the establishment of this model lays the groundwork for future experiments including CD8+ T lymphocyte depletion and B lymphocyte depletion studies in CMV-seropositive macaques to dissect the contribution of different arms of the adaptive immune system involved in protection against non-primary cCMV. Taken together, our data reinforces the utility of the rhesus macaque model in furthering knowledge about immune determinants of cCMV protection, needed for rational vaccine design.

## Methods

### Animals and study design

A total of eight CMV-seropositive Indian-origin first trimester of gestation rhesus macaques were enrolled in the study from the specific pathogen free colony at the Tulane National Primate Research Center (TNPRC) and the New England Primate Research Center (NEPRC) (**Table 1**). Gestational age at enrollment was estimated by ultrasound measurement of gestational sac diameter and/or crown-rump length. Subsequently, the gestational age was monitored weekly by ultrasound measurement of biparietal diameter and femur length. At Caesarian section (C-section), measurements of occipitofrontal diameter, head circumference, and abdominal circumference were recorded.

All eight dams were subjected to *in vivo* CD4+ T lymphocyte depletion by intravenous (IV) administration of 50 mg/kg rhesus-recombinant anti-CD4 antibody (Clone CD4R1; Nonhuman Primate Reagent Resource) at mean 52.3 gestation days (range 49-59). The rhesus IgG1 recombinant Anti-CD4 [CD4R1] monoclonal antibody was engineered and produced by the Nonhuman Primate Reagent Resource (NIH Nonhuman Primate Reagent Resource Cat# PR-0407, RRID:AB_2716322).

Five of eight dams were reinfected with RhCMV one week after administration of the CD4-depleting antibody (CMV-seropositive reinfection group), whereas the remaining three dams that received the CD4-depleting antibody were not reinfected and served as a control group for reactivation following CD4+ T lymphocyte depletion (CMV-seropositive Controls). Data from six dams with CD4-depletion and a primary infection with RhCMV served as historical controls (**Table 1**)(24, 25).

One week after administration of the anti-CD4 antibody, the CMV-seropositive reinfection group were inoculated IV with 2×10^6^ TCID_50_ of RhCMV 180.92 (n=2) or 1×10^6^ pfu each of RhCMV UCD52 and FL-RhCMVΔRh13.1/SIV*gag* (n=3; **Table 1**). Maternal blood, saliva, urine, amniotic fluid were collected preceding anti-CD4+ T lymphocyte depletion and virus inoculation. Following RhCMV inoculation the animals were sampled weekly for all sample types. Three animals that received the RhCMV UCD52 and FL-RhCMVΔRh13.1/SIV*gag* were in addition sampled for blood, urine and saliva every 1-3 days within the first 7 days post infection to study the acute phase of infection. All animals were subsequently followed with weekly sampling until term or C-section, outlined in **Table 1**.

### Virus stocks for reinfection

Full Length-RhCMV68-1-ΔRh13.1/SIV*gag* (RhCMVΔRh13.1/SIV*gag*) virus was generated and characterized as previously described (26) was administered intravenously at a single dose of 1×10^6^ pfu. RhCMV strain UCD52 virus stock grown in TeloRF cells was provided by Dr. Peter Barry (41) and administered intravenously at a single dose of 1×10^6^ pfu. Both virus strains were inoculated in to separate arms of three animals. RhCMV 180.92 virus stock grown in primary rhesus fibroblasts as previously described (27, 28) was administered at a single dose of 2×10^6^ TCID_50_ intravenously to two animals.

### Sample collection and processing

Maternal PBMC’s were isolated by ficoll separation after collecting plasma. All PBMC were cryopreserved using 90%FBS/10% DMSO. Amniotic fluid was spun to remove debris prior to storage in aliquots at-20^0^C. Saliva and urine sample supernatants were concentrated using Ultracel YM-30 (Amicon/Milipore) and subsequently aliquoted for storage at-20^0^C for DNA extraction. At C-section, the placenta and fetal tissues were harvested and processed for lymphocyte isolation, snap frozen for DNA extraction for PCR and placed in Z-fix for paraffin blocks.

DNA was extracted from urine with the QIAmp RNA mini kit (Qiagen, Velencia, CA); from amniotic fluid, plasma, and saliva with the QIAmp DNA mini kit (Qiagen, Velencia, CA); and from 10-25mgm of snap-frozen tissue using the DNeasy Blood and Tissue kit (Qiagen) as previously described (21, 24).

### Viral quantitation by real time PCR

Absolute quantification of RhCMV DNA in tissues and maternal fluids were performed as previously described (24, 42). Briefly, for RhCMV DNA PCR, the primers/probe targeting the noncoding exon 1 region of the immediate early gene were used. The Forward primer 5’-GTTTAGGGAACCGCCATTCTG-3’, Reverse primer 5’-GTATCCGCGTTCCAATGCA-3’, and probe 5’-FAM-TCCAGCCTCCATAGCCGGGAAGG-TAMRA-3’ were used in a 25μL reaction with Supermix Platinum™ Quantitative PCR SuperMix-UDG (Invitrogen). The reaction was performed in a 96-well format for real time quantification on Applied Biosystems 7900HT Fast Real-Time PCR System. A standard curve generated from amplification of 10^5^-10^0^ copies of plasmid standard containing the RhCMV IE target sequence diluted in genomic DNA from CMV-seronegative rhesus macaques was used for absolute quantitation of RhCMV DNA. Real time PCR was performed in 6-12 replicates and at least 2 positive replicates were required to be reported as a positive result. Data of plasma and amniotic fluid were reported as RhCMV DNA copies per mL of sample while saliva, urine, and tissues were reported as copies per microg of input DNA.

For absolute quantification of SIV*gag* DNA in the three animals which received FL-RhCMVΔRh13.1/SIV*gag* both a nested and real-time PCR protocol was performed. A plasmid carrying SIV*gag* sequence was synthesized (Intregrated DNA Technology, Iowa) and used as standard in a 25uL reaction with Platinum Taq DNA polymerase (cat# 10966034 Invitrogen) mastermix containing 0.012% Tween 20, 0.006% gelatin, 4.5mM MgCl_2_, 300μM dNTPs, 10% PCR II buffer, 300nM Forward 5’-CAACTACGTCAACCTGCCACTGTC-3’, 300nM Reverse 5’-TCCAACGCAGTTCAGCATCTGG-3’, 200nM Probe 5’-FAM-CCGAGAACCCTGAACGCTTGGGTCAAGC-3BHQ-3’. This was performed in 96 well plate format on Applied Biosystems 7900HT Fast Real-Time PCR System at 95°C for 2 minutes and cycled for 45 cycles at 95°C for 15 seconds, and 60°C for 1 minute.

### Viral sequencing and analysis

Ultradeep sequencing of RhCMV DNA in the amniotic fluid of KK24 generally followed amplicon based methodologies previously established for sequencing human CMV genomes (43) as applied to RhCMV (44). These studies demonstrated that the error rates observed in amplicon based sequencing and direct sequencing of BAC clones for both viral species are very low and produce nearly identical results. Thus, errors resulting from the amplicon-based workflow plus sequencing are very similar in type and rate to those introduced by direct sequencing and applicable to differentiating viral strains in a sample. As the amount of viral DNA in the KK24 amniotic fluid was limited, MDA (Repli-g, Qiagen) was initially used to increase the overall amounts of DNA for the sequencing workflow. The resulting DNA was repurified and subjected to plate PCR using primer pairs that span the parental RhCMV genome and the SIV*gag* insert, fragmented and then ligated to Ion Express barcodes (Life Technologies). Products were pooled with final processing performed on an Ion Chef (Life Technologies) followed by sequencing on an Ion Proton Sequencer (Life Technologies).

Whole-genome sequencing data was mapped against the RhCMV UCD52 and FL-RhCMVΔRh13.1/SIV*gag* reference assemblies using NextGenMap v.0.5.15 (45). Thereby, the human (hg38) and rhesus (rheMac10) genomes (downloaded from NCBI GenBank using accession numbers GCA_000001405.29 and GCA_003339765.3, respectively) were included as decoys to remove any potential contamination. To distinguish between the RhCMV strains, uniquely mapping regions were identified using SAMtools v.1.12.0 (46).

### Cytokine and Chemokine analysis Luminex

To analyze peripheral soluble cytokines and chemokines, a luminex assay was performed with Cytokine & Chemokine 30-Plex NHP ProcartaPlex™ Panel (Invitrogen, EPX300-40044-901). The analytes in this panel are BLC (CXCL13); Eotaxin (CCL11); G-CSF (CSF-3); GM-CSF; IFN alpha; IFN gamma; IL-1 beta; IL-10; IL-12p70; IL-13; IL-15; IL-17A (CTLA-8); IL-18; IL-1RA; IL-2; IL-23; IL-4; IL-5; IL-6; IL-7; IL-8 (CXCL8); IP-10 (CXCL10); I-TAC (CXCL11); MCP-1 (CCL2); MIG (CXCL9); MIP-1 alpha (CCL3); MIP-1 beta (CCL4); sCD40L; SDF-1 alpha (CXCL12a); TNF alpha. Plasma was thawed on ice and manufacturers instructions were followed to prepare a 96-well plate with samples performed in duplicates and read on a Bio-Plex® 200 System (Bio-Rad Laboratories, Hercules, CA). Results were calculated using Bio-Plex Manager™ Software v6.2 (Bio-Rad) and the mean concentration of each analyte was plotted.

### Immunophenotyping and intracellular cytokine staining (ICS) assays

CD4+ T lymphocyte depletion kinetics were monitored by flow cytometric evaluation of absolute counts. Briefly, 50 μL whole blood was stained with an 8-color panel of FITC-CD3, PerCP-CD45, APC-CD4, V500-CD8, PE-Cy7-CD95, APC-CY7-CD20, BV421-CCR7. A FMO was performed for CCR7 which was first stained alone for 15 minutes and then with the remaining cocktail for an additional 15 minutes. Red blood cells were lysed using BD Lysing buffer for 15-20 minutes and subsequently aquired on a BD FACSverse.

A 13-color flow cytometry panel was used for immunophenotyping of the acute reponses following RhCMV infection. PBMCs were stained with the following antibodies: FITC-Ki67, PCP-Cy5.5-TCRgd, APC-KIR2D, AL700-Granzyme B, APC-CY7-CD3, PacBlue-CD20, BV510-live/dead, BV605-CD14, BV650-CD8, BV711-CD16, PE-CD169, PE-CF594-HLA-DR, PE-CY7-NKG2A (for additional details regarding the antibodies, see **Table S2**). These data were aquired on the BD LSRFortessa™ and analyzed using Flowjo v9.9 (Ashland, Oregon).

Antigen-specific T lymphocyte respones were assesed by intracellular cytokine staining for RhCMV-specific and SIV Gag-specific responses. Cryopreseved PBMCs were thawed and stimulated for 12-18 hours with RhCMV IE1, IE2, pp65 and SIV Gag peptide pools. Briefly, the RhCMV peptide pool and the SIV Gag peptide pool consists of pools of 15-amino acid peptides each, overlapping by 11 amnio acids and spanning the entire protein. Peptides were used at a concentration of 1μg/mL of individual peptides for stimulation and DMSO concentrations were kept <0.5%. After 1 hour, monensin 2μM/mL (cat# 554724, BD) and bredfeldin A 1μL/mL (cat#555029, BD) were added along with CD107a-FITC and CD107b-FITC for the remaining period of stimulation. After stimulation, cells were washed and stained sequentially with the following: AQUA Live/dead dye, surface, and intracellular cytokine staining antibodies PCP-Cy5.5-CD4, APC-CD69, AL700-TNFα, APC-CY7-CD3, BV421-Granzyme B, BV605-IL-2, BV650-CD8, BV711-CD95, PE-CCR5, PE-CF594-CD28, PE-CY7-IFNγ using the BD fix/perm kit(BD Cat# 554714) and Brilliant Stain buffer(BD Cat# 563794). These data were aquired on the BD LSRFortessa™ and analyzed using Flowjo v9.9 (Ashland, Oregon). Boolean analyses of polyfunctional responses were performed using the SPICE 6 software (47).

### Antibody assays for RhCMV gB and SIV Gag

#### RhCMV gB

DNA plasmid expressing RhCMV gB was transected into 293F/293i cells. Protein was purified with nickel beads and quantitated on a NanoDrop.

#### IgG ELISA

ELISAs were performed as previously described (24). Briefly 384-well high protein binding plates (Corning 3700) were coated overnight at 4^0^C with either 30 ng RhCMV gB produced as described above or SIVmac251 pr55 Gag recombinant protein (NIH AIDS Reagent Program cat# 13384). Serially diluted plasma samples were added to the coated ELISA plates, incubated for 2 hours after which the plates were washed, and then incubated with the secondary anti-monkey IgG HRP-conjugated antibody followed by addition of substrate as previously described (24). ELISA titers were reported as the serum dilution yielding 50% maximum absorbance (Effective Dose ED_50_). ED_50_ was determined using non-linear regression in GraphPad Prism v9.5. Data are reported as log^10^ED_50_.

#### Antibody neutralization assay

Telo-RF cells were seeded in a 384-well plate and incubated for 1 day at 37^0^C at 5% CO2. The next day, serial dilutions (1:10-1:21,870) of heat-inactivated rhesus plasma were incubated with 1 PFU of RhCMV 180.92 per cell. Infected cells were then fixed for 20 minutes at-20^0^C with 1:1 methanol/acetone, rehydrated in PBS three times for 5 minutes and stained with 1 mg/mL mouse anti-RhCMV-IE1 monoclonal antibody provided by Dr. Klaus Früh (Oregon Health and Science University, Portland, OR) followed by a 1:1000 dilution of anti-mouse IgG-Alexa Fluor 488 antibody. Nuclei were stained with DAPI for 5 minutes (Pierce) and imaged using the CellInsight CX5 High-Content Screening (HCS) platform. The 50% neutralization titer (NT50) was determined by comparing the dilution that resulted in a 50% reduction in fluorescence signal to control wells infected with virus only.

### Statistics

Unpaired and paired parametric and non-parametric t-tests were performed in GraphPad Prism version 8.4.0. (San Diego, CA). P-values <0.05 were considered significant. For viral load the area under the curve (AUC) was calculated between day 0 and 99 days post infection for inter-group comparison. Kaplan-Meier survival curves were calculated in Graphpad Prism for transmission events and fetal survival and statistical comparisons performed with the Log-rank (Mantel-Cox) test.

## Acknowledgements

This work was supported by NIH grants DP2HD075699, P01AI129859, and P51OD011104. The Anti-CD4 [CDR1] antibody used in this study was provided by the NIH Nonhuman Primate Reagent Resource (P40 OD028116). The authors gratefully acknowledge the Veterinary Medicine staff at the Tulane National Primate Reseach Center (TNPRC) for care of the animals, TNPRC Flow Cytometry Core for acquisition of flow cytometry data and TNPRC Pathogen Detection and Quantification Core for real time PCR runs and Luminex assays.

## References

1. Cheeran MC, Lokensgard JR, Schleiss MR. Neuropathogenesis of congenital cytomegalovirus infection: disease mechanisms and prospects for intervention. Clin Microbiol Rev. 2009;22(1):99–126.

2. Cannon MJ, Schmid DS, Hyde TB. Review of cytomegalovirus seroprevalence and demographic characteristics associated with infection. Rev Med Virol. 2010;20(4):202–13.

3. Ahlfors K, Ivarsson SA, Harris S. Secondary maternal cytomegalovirus infection--A significant cause of congenital disease. Pediatrics. 2001;107(5):1227–8.

4. Boppana SB, Rivera LB, Fowler KB, Mach M, Britt WJ. Intrauterine transmission of cytomegalovirus to infants of women with preconceptional immunity. The New England journal of medicine. 2001;344(18):1366–71.

5. Ross SA, Arora N, Novak Z, Fowler KB, Britt WJ, Boppana SB. Cytomegalovirus reinfections in healthy seroimmune women. J Infect Dis. 2010;201(3):386–9.

6. Yamamoto AY, Mussi-Pinhata MM, Boppana SB, Novak Z, Wagatsuma VM, Oliveira Pde F, et al. Human cytomegalovirus reinfection is associated with intrauterine transmission in a highly cytomegalovirus-immune maternal population. Am J Obstet Gynecol. 2010;202(3):297 e1-8.

7. Wang C, Zhang X, Bialek S, Cannon MJ. Attribution of congenital cytomegalovirus infection to primary versus non-primary maternal infection. Clin Infect Dis. 2011;52(2):e11–3.

8. Plotkin SA, Boppana SB. Vaccination against the human cytomegalovirus. Vaccine. 2019;37(50):7437–42.

9. Britt WJ. Congenital Human Cytomegalovirus Infection and the Enigma of Maternal Immunity. J Virol. 2017;91(15).

10. Kenneson A, Cannon MJ. Review and meta-analysis of the epidemiology of congenital cytomegalovirus (CMV) infection. Rev Med Virol. 2007;17(4):253–76.

11. Leruez-Ville M, Magny JF, Couderc S, Pichon C, Parodi M, Bussieres L, et al. Risk Factors for Congenital Cytomegalovirus Infection Following Primary and Nonprimary Maternal Infection: A Prospective Neonatal Screening Study Using Polymerase Chain Reaction in Saliva. Clin Infect Dis. 2017;65(3):398–404.

12. Simonazzi G, Curti A, Cervi F, Gabrielli L, Contoli M, Capretti MG, et al. Perinatal Outcomes of Non-Primary Maternal Cytomegalovirus Infection: A 15-Year Experience. Fetal Diagnosis and Therapy. 2018;43(2):138–42.

13. Mussi-Pinhata MM, Yamamoto AY, Aragon DC, Duarte G, Fowler KB, Boppana S, et al. Seroconversion for Cytomegalovirus Infection During Pregnancy and Fetal Infection in a Highly Seropositive Population: “The BraCHS Study”. J Infect Dis. 2018;218(8):1200–4.

14. Giannattasio A, Di Costanzo P, De Matteis A, Milite P, De Martino D, Bucci L, et al. Outcomes of congenital cytomegalovirus disease following maternal primary and non-primary infection. J Clin Virol. 2017;96:32–6.

15. Ornoy A, Diav-Citrin O. Fetal effects of primary and secondary cytomegalovirus infection in pregnancy. Reprod Toxicol. 2006;21(4):399–409.

16. Yamamoto AY, Anastasio ART, Massuda ET, Isaac ML, Manfredi AKS, Cavalcante JMS, et al. Contribution of Congenital Cytomegalovirus Infection to Permanent Hearing Loss in a Highly Seropositive Population: The Brazilian Cytomegalovirus Hearing and Maternal Secondary Infection Study. Clin Infect Dis. 2020;70(7):1379–84.

17. Manicklal S, Emery VC, Lazzarotto T, Boppana SB, Gupta RK. The “Silent” Global Burden of Congenital Cytomegalovirus. Clinical Microbiology Reviews. 2013;26(1):86–102.

18. Deere JD, Barry PA. Using the nonhuman primate model of HCMV to guide vaccine development. Viruses. 2014;6(4):1483–501.

19. Itell HL, Kaur A, Deere JD, Barry PA, Permar SR. Rhesus monkeys for a nonhuman primate model of cytomegalovirus infections. Curr Opin Virol. 2017;25:126–33.

20. Malouli D, Nakayasu ES, Viswanathan K, Camp DG, 2nd, Chang WL, Barry PA, et al. Reevaluation of the coding potential and proteomic analysis of the BAC-derived rhesus cytomegalovirus strain 68-1. J Virol. 2012;86(17):8959–73.

21. Antoine P, Varner V, Carville A, Connole M, Marchant A, Kaur A. Postnatal acquisition of primary rhesus cytomegalovirus infection is associated with prolonged virus shedding and impaired CD4+ T lymphocyte function. J Infect Dis. 2014;210(7):1090–9.

22. Kaur A, Itell HL, Ehlinger EP, Varner V, Gantt S, Permar SR. Natural history of postnatal rhesus cytomegalovirus shedding by dams and acquisition by infant rhesus monkeys. PLoS One. 2018;13(10):e0206330.

23. Vogel P, Weigler BJ, Kerr H, Hendrickx AG, Barry PA. Seroepidemiologic studies of cytomegalovirus infection in a breeding population of rhesus macaques. Lab Anim Sci. 1994;44(1):25–30.

24. Bialas KM, Tanaka T, Tran D, Varner V, Cisneros De La Rosa E, Chiuppesi F, et al. Maternal CD4+ T cells protect against severe congenital cytomegalovirus disease in a novel nonhuman primate model of placental cytomegalovirus transmission. Proceedings of the National Academy of Sciences of the United States of America. 2015;112(44):13645–50.

25. Nelson CS, Cruz DV, Tran D, Bialas KM, Stamper L, Wu H, et al. Preexisting antibodies can protect against congenital cytomegalovirus infection in monkeys. JCI Insight. 2017;2(13).

26. Taher H, Mahyari E, Kreklywich C, Uebelhoer LS, McArdle MR, Mostrom MJ, et al. In vitro and in vivo characterization of a recombinant rhesus cytomegalovirus containing a complete genome. PLoS Pathog. 2020;16(11):e1008666.

27. Assaf BT, Mansfield KG, Strelow L, Westmoreland SV, Barry PA, Kaur A. Limited dissemination and shedding of the UL128 complex-intact, UL/b’-defective rhesus cytomegalovirus strain 180.92. J Virol. 2014;88(16):9310–20.

28. Rivailler P, Kaur A, Johnson RP, Wang F. Genomic sequence of rhesus cytomegalovirus 180.92: insights into the coding potential of rhesus cytomegalovirus. J Virol. 2006;80(8):4179–82.

29. Vera Cruz D, Nelson CS, Tran D, Barry PA, Kaur A, Koelle K, et al. Intrahost cytomegalovirus population genetics following antibody pretreatment in a monkey model of congenital transmission. PLoS Pathog. 2020;16(2):e1007968.

30. Hansen SG, Ford JC, Lewis MS, Ventura AB, Hughes CM, Coyne-Johnson L, et al. Profound early control of highly pathogenic SIV by an effector memory T-cell vaccine. Nature. 2011;473(7348):523–7.

31. Hansen SG, Vieville C, Whizin N, Coyne-Johnson L, Siess DC, Drummond DD, et al. Effector memory T cell responses are associated with protection of rhesus monkeys from mucosal simian immunodeficiency virus challenge. Nat Med. 2009;15(3):293–9.

32. Hansen SG, Powers CJ, Richards R, Ventura AB, Ford JC, Siess D, et al. Evasion of CD8+ T cells is critical for superinfection by cytomegalovirus. Science. 2010;328(5974):102–6.

33. Ross SA, Novak Z, Pati S, Patro RK, Blumenthal J, Danthuluri VR, et al. Mixed infection and strain diversity in congenital cytomegalovirus infection. J Infect Dis. 2011;204(7):1003–7.

34. Nyland TG, Hill DE, Hendrickx AG, Farver TB, McGahan JP, Henrickson R, et al. Ultrasonic assessment of fetal growth in the nonhuman primate (Macaca mulatta). J Clin Ultrasound. 1984;12(7):387–95.

35. Hyde TB, Schmid DS, Cannon MJ. Cytomegalovirus seroconversion rates and risk factors: implications for congenital CMV. Rev Med Virol. 2010;20(5):311–26.

36. Barbosa NG, Yamamoto AY, Duarte G, Aragon DC, Fowler KB, Boppana S, et al. Cytomegalovirus Shedding in Seropositive Pregnant Women From a High-Seroprevalence Population: The Brazilian Cytomegalovirus Hearing and Maternal Secondary Infection Study. Clin Infect Dis. 2018;67(5):743–50.

37. Renzette N, Bhattacharjee B, Jensen JD, Gibson L, Kowalik TF. Extensive Genome-Wide Variability of Human Cytomegalovirus in Congenitally Infected Infants. PLoS Pathogens. 2011;7(5).

38. Pokalyuk C, Renzette N, Irwin KK, Pfeifer SP, Gibson L, Britt WJ, et al. Characterizing human cytomegalovirus reinfection in congenitally infected infants: an evolutionary perspective. Mol Ecol. 2017;26(7):1980–90.

39. Novak Z, Ross SA, Patro RK, Pati SK, Kumbla RA, Brice S, et al. Cytomegalovirus strain diversity in seropositive women. Journal of clinical microbiology. 2008;46(3):882–6.

40. Ezekian B, Schroder PM, Mulvihill MS, Barbas A, Collins B, Freischlag K, et al. Pretransplant Desensitization with Costimulation Blockade and Proteasome Inhibitor Reduces DSA and Delays Antibody-Mediated Rejection in Highly Sensitized Nonhuman Primate Kidney Transplant Recipients. J Am Soc Nephrol. 2019;30(12):2399–411.

41. Oxford KL, Strelow L, Yue Y, Chang WL, Schmidt KA, Diamond DJ, et al. Open reading frames carried on UL/b’ are implicated in shedding and horizontal transmission of rhesus cytomegalovirus in rhesus monkeys. J Virol. 2011;85(10):5105–14.

42. Kaur A, Kassis N, Hale CL, Simon M, Elliott M, Gomez-Yafal A, et al. Direct relationship between suppression of virus-specific immunity and emergence of cytomegalovirus disease in simian AIDS. J Virol. 2003;77(10):5749–58.

43. Renzette N, Pokalyuk C, Gibson L, Bhattacharjee B, Schleiss MR, Hamprecht K, et al. Limits and patterns of cytomegalovirus genomic diversity in humans. Proceedings of the National Academy of Sciences of the United States of America. 2015;112(30):E4120–8.

44. Deere JD, Chang WL, Castillo LD, Schmidt KA, Kieu HT, Renzette N, et al. Utilizing a TLR5-Adjuvanted Cytomegalovirus as a Lentiviral Vaccine in the Nonhuman Primate Model for AIDS. PLoS One. 2016;11(5):e0155629.

45. Sedlazeck FJ, Rescheneder P, von Haeseler A. NextGenMap: fast and accurate read mapping in highly polymorphic genomes. Bioinformatics. 2013;29(21):2790–1.

46. Danecek P, Bonfield JK, Liddle J, Marshall J, Ohan V, Pollard MO, et al. Twelve years of SAMtools and BCFtools. Gigascience. 2021;10(2).

47. Roederer M, Nozzi JL, Nason MC. SPICE: exploration and analysis of post-cytometric complex multivariate datasets. Cytometry A. 2011;79(2):167–74.

