## Supplemental Figures and Tables for "Protective effect of pre-existing natural immunity in a nonhuman primate reinfection model of congenital cytomegalovirus infection"

Mmostrom et al.

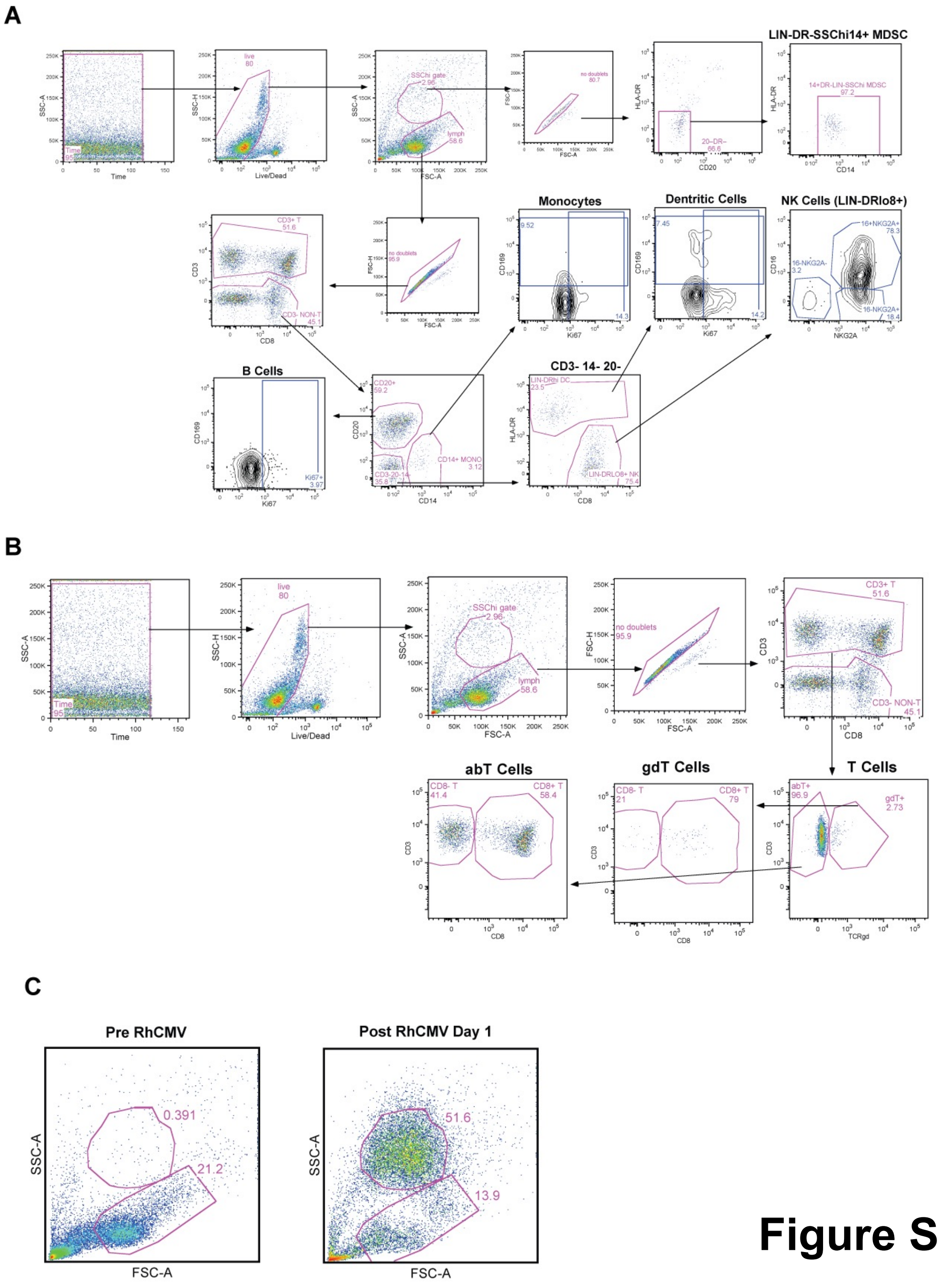


### Figure S1

**S1 Fig. Gating strategy for PBMC immunophenotyping analysis in acute RhCMV reinfection**

(A) Gating strategy used for the innate immune cell compartment. (B) The gating strategy used to identify

T cell subsets. (C) Representative plots of side scatter high (SSChi) population following RhCMV reinfection in CMV-seropositive rhesus macaque dams.

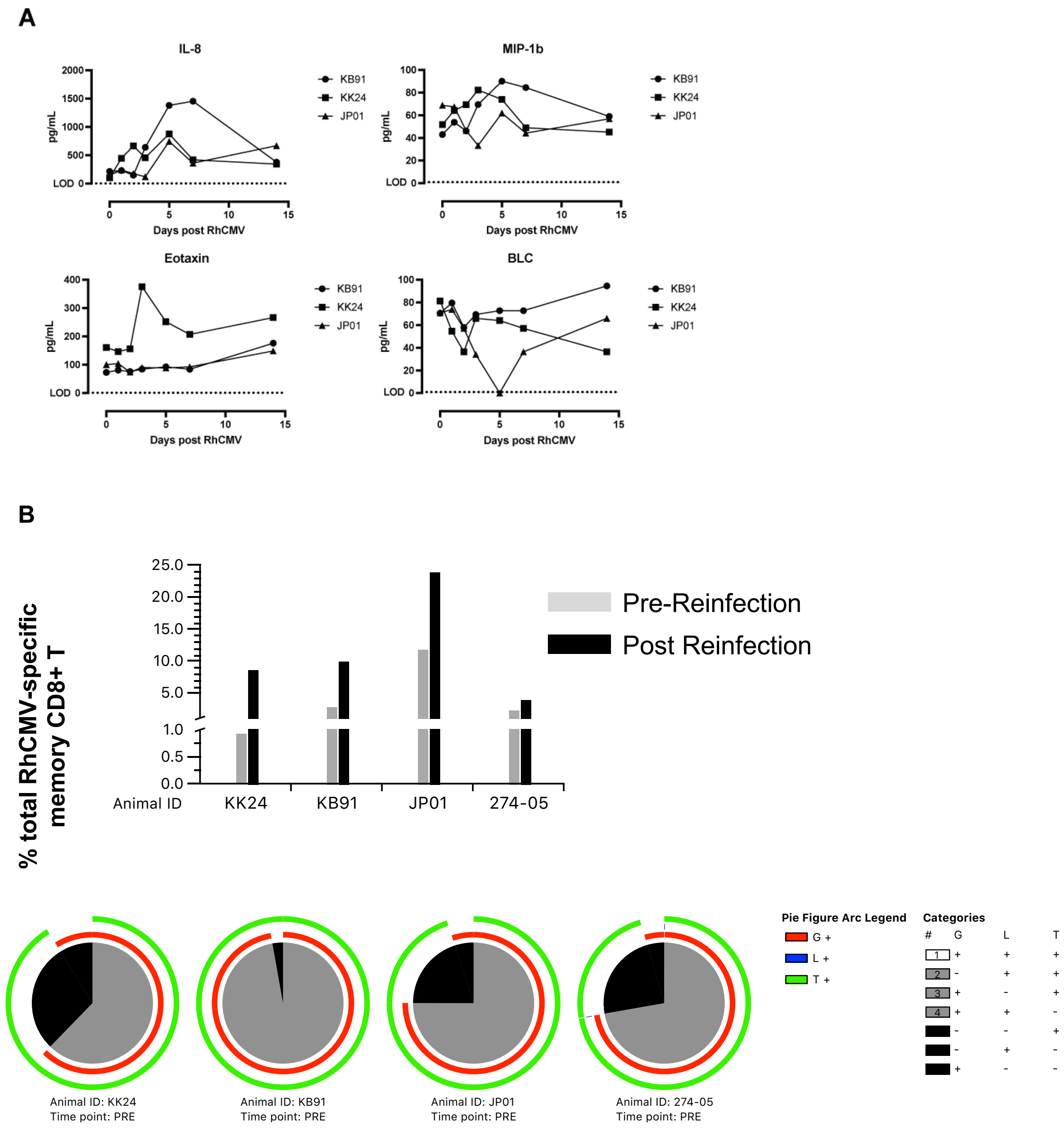


### Figure S2

**S2 Fig. Innate and adaptive immune responses in CMV-seropositive reinfected dams.**

(A) Plasma IL-8, MIP-1b, Eotaxin, and B-lymphocyte chemoattractant (BLC) levels in three RhCMV reinfected dams in the first two weeks post RhCMV reinfection. Data generated using a nonhuman primate (NHP) 30-plex Luminex assay. Limit of detection (LOD) of the lot# is shown as a stippled line at the bottom of the y-axis. (B) Total memory RhCMV IE-specific CD8+ T lymphocyte responses at pre- and post reinfection time-points in four dams shown in the top panel. Post reinfection time-points varied between week 8 (KK24, KB91, JP01) and week 10 (274-05) post reinfection. Bottom panel showing pie charts depicting proportion of 3-functional, 2-functional and mono-functional RhCMV IE-specific CD8+ T lymphocyte responses prior to reinfection in the four dams.

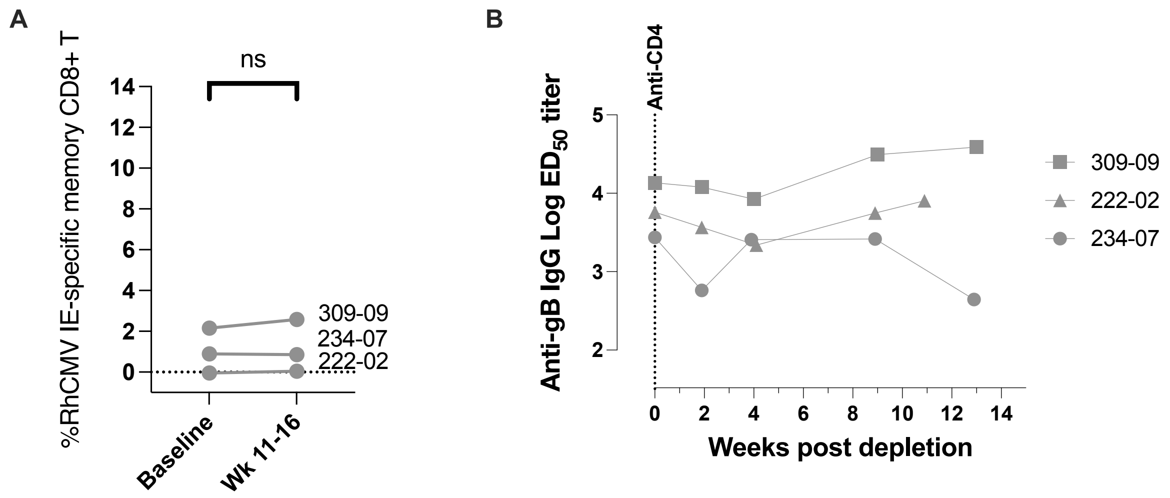


### Figure S3

**S3 Fig. RhCMV-specific adaptive immune responses in CD4+ T lymphocyte depleted CMV-seropositive control macaques.**

(A) Memory CD8+ T lymphocyte responses to RhCMV IE protein in CMV-seropositive controls. (B) Kinetics of RhCMV gB-specific binding antibodies in CMV-seropositive controls.

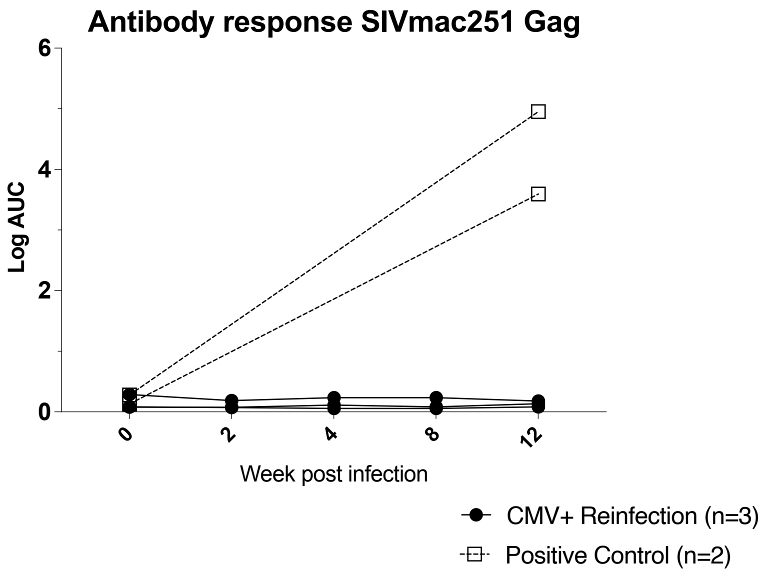


### Figure S4

**S4 Fig. Gag-specific binding antibody assay in CD4+ T lymphocyte depleted CMV-seropositive rhesus macaque dams that received RhCMV FL-RhCMVΔRh13.1/SIV*gag*.**

Positive controls in this assay are SHIV-infected rhesus macaques (open symbol) were used for comparison of responses in CMV-seropositive reinfected animals (closed symbol).

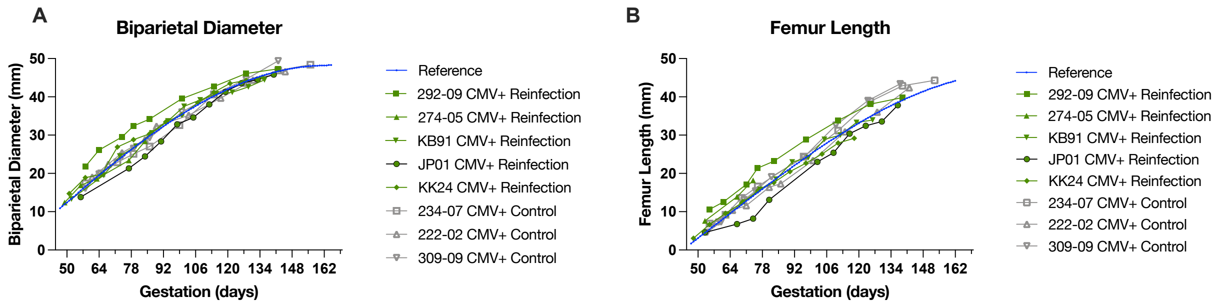


### Figure S5

**S5 Fig. Ultrasound measurements over time of Biparietal Diameter (BPD) and Femur Length (FL)**

(A) BPD of CMV-seropositive reinfected (green) and CMV-seropositive controls (grey) fetuses compared to reference values (34). (B) FL of CMV-seropositive reinfected (green) and CMV-seropositive control (grey) fetuses compared to reference values (34).

| Group | | CMV-seropostive Reinfection | | | CMV-seronegative Primary Infection | | | | | |
|---|---|---|---|---|---|---|---|---|---|---|
| Animal ID | | KB91 | KK24 | JP01 | HD79 | GM04 | 145-97 | 369-09 | 174-97 | 274-98 |
| Amniotic fluid | Peak mean RhCMV DNA copies/mL | - | 57 | - | 521 | 482 | 580 | 372 | 249 | 49 |
| Placental tissues | 1' Placental maternal side | - | - | - | 1438 | 100 | NS | 24228 | 2477 | 245 |
| | 1' Placental fetal side | - | - | - | NS | NS | NS | NS | NS | NS |
| | 2' Placental maternal side | NS | - | NS | 630 | 28 | NS | NS | NS | NS |
| | 2' Placental fetal side | NS | - | NS | 424 | NS | NS | NS | NS | NS |
| | Decidua | - | - | - | NS | NS | NS | NS | NS | NS |
| | Umbilical cord | - | - | - | 7 | NS | NS | 2699 | 3920 | NS |
| | Amniotic membrane | - | - | - | 1902 | 2746 | NS | NS | NS | NS |
| Fetal tissues | FAT | - | - | - | NS | NS | NS | NS | NS | NS |
| | Pheripheral Lymph Node | - | - | - | NS | NS | 601 | NS | NS | NS |
| | Mesenteric Lymph Node | | NS | | NS | NS | - | NS | NS | NS |
| | Thymus | - | - | - | NS | NS | NS | NS | NS | NS |
| | Jejunum | - | - | - | - | NS | - | NS | 1220 | NS |
| | Colon | - | - | - | NS | NS | - | NS | NS | NS |
| | Ileum | - | - | - | NS | NS | NS | NS | NS | NS |
| | Liver | - | - | - | - | NS | 1 | 709 | 1849 | NS |
| | Spleen | - | - | - | - | NS | 1,611 | NS | 880 | NS |
| | Kidney | - | - | - | NS | NS | 78,643 | NS | NS | NS |
| | Cervical SC | NS | NS | NS | NS | NS | 21 | NS | NS | NS |
| | Brain (frontal) | - | - | - | - | NS | - | NS | 696 | NS |
| | Brain(mid) | - | - | - | - | NS | NS | NS | NS | NS |
| | Brain (occipipital) | - | - | - | - | NS | NS | NS | 595 | NS |
| | Brain stem | NS | NS | NS | NS | NS | - | NS | NS | NS |
| | Cerebellum | - | - | - | - | NS | NS | NS | NS | NS |
| | Ear cochlea (left or right) | - | - | - | 313 | NS | NS | NS | NS | NS |
| | Paratoid Salivary gland | NS | NS | NS | NS | NS | 5,405 | NS | NS | NS |
| | Submandibulary Salivary gland | NS | NS | NS | NS | NS | 60 | NS | NS | NS |
| | Heart/Left Ventricle | - | - | - | - | NS | 665 | NS | 2497 | NS |
| | Lung | - | - | - | - | NS | 179 | NS | 1114 | NS |
| Fetal Outcome | C-section/Live Birth/Spontanious Abortion | C-section | C-section | C-section | Spontaneous abortion | Spontaneous abortion | Live birth | Spontaneous abortion | Spontaneous abortion | Spontaneous abortion |

**S1 Table. RhCMV DNA PCR in placental and fetal tissues of CD4+ T lymphocyte-depleted dams.** NS = No sample. RhCMV DNA copy numbers shown as a heatmap; increasing values in shades of red. RhCMV DNA expressed as copies per µg input DNA in tissues and copies/ml in amniotic fluid.

| Target | Fluorochrome | Company | Clone | Panel | Surface/Intracellular |
|---|---|---|---|---|---|
| Live/Dead™ | BV510 | Invitrogen | Aqua | Phenotyping | Surface |
| CD3 | APC-Cy7 | BD | SP34-2 | Phenotyping/ICS | Surface |
| CD4 | PerCP-Cy5.5 | BD | L200 | ICS | Surface |
| CD8 | BV650 | BD | SK1 | Phenotyping/ICS | Surface |
| CD14 | BV605 | BD | M5E2 | Phenotyping | Surface |
| CD16 | BV711 | Biolegend | 3G8 | Phenotyping | Surface |
| CD20 | PacBlue | Biolegend | 2H7 | Phenotyping | Surface |
| CD28 | PE-CF594 | BD | CD28.2 | ICS | Surface |
| CD69 | APC | BioLegend | FN50 | ICS | Intracellular |
| CD95 | BV711 | BD | DX2 | ICS | Surface |
| CD107a | FITC | BD | H4A3 | ICS | Surface (1h into stimulation) |
| CD107b | FITC | BD | H4B4 | ICS | Surface (1h into stimulation) |
| CD169 | PE | Biolegend | 7-239 | Phenotyping | Surface |
| CD195 (CCR5) | PE | BD | 3A9 | ICS | Surface |
| Ki-67 | FITC | BD | B56 | Phenotyping | Intracellular |
| TCR γδ | PCP-Cy5.5 | BioLegend | B1 | Phenotyping | Surface |
| KIR2D | APC | Miltenyi | NKVFS1 | Phenotyping | Surface |
| Granzyme B | AL700 | BD | GB11 | Phenotyping | Intracellular |
| Granzyme B | BV421 | BD | GB11 | ICS | Intracellular |
| HLA-DR | PE-CF594 | BD | G46-6 | Phenotyping | Surface |
| NKG2A | PE-Cy7 | Beckman/Coulter | Z199 | Phenotyping | Surface |
| IFNγ | PE-Cy7 | BD | B27 | ICS | Intracellular |
| IL-2 | BV605 | BD | MQ1-17H12 | ICS | Intracellular |
| TNFα | AL700 | BD | Mab11 | ICS | Intracellular |

**S2 Table. Antibodies for phenotyping and intracellular cytokine staining.**
